# Runx factors launch T-cell and innate lymphoid programs via direct and gene network-based mechanisms

**DOI:** 10.1101/2022.11.18.517146

**Authors:** Boyoung Shin, Wen Zhou, Jue Wang, Fan Gao, Ellen V. Rothenberg

## Abstract

Runx factors are essential for lineage specification of various hematopoietic cells, including T lymphocytes. However, they regulate context-specific genes and occupy distinct genomic regions in different cell types. Here, we show that dynamic Runx binding shifts in early T-cell development are mostly not restricted by local chromatin state but regulated by Runx dosage and functional partners. Runx co-factors compete to recruit a limited pool of Runx factors in early T-progenitors, and a modest increase in Runx protein availability at pre-commitment stages causes premature Runx occupancy at post-commitment binding sites. This results in striking T-lineage developmental acceleration by selectively activating T-identity and innate lymphoid cell programs. These are collectively regulated by Runx together with other, Runx-induced transcription factors that co-occupy Runx target genes and propagate gene network changes.

## Introduction

Runx family transcription factors (Runx1, Runx2, Runx3, and their cofactor CBFβ) are essential for T cell development from the earliest steps in the lineage, playing partially redundant roles^1–6^. However, they are also vital for the establishment of hematopoietic stem cells in early embryos^7^ and for generation of B cells and megakaryocytes throughout life^8–11^, quite different programs. Early intrathymic T cell development itself spans distinct regulatory contexts where Runx factors can implement stage-unique or continuing functional inputs at different stages. Runx target motifs are consistently highly enriched around open chromatin sites and lineage-specific transcription factor (TF) binding sites in multiple hematopoietic lineages^12–19^, suggesting a common contribution to active enhancers generally. However, we have recently shown that Runx factors occupy different genomic sites at different stages of early T cell development, and that this results in their regulation of different target genes^1^. Thus, key questions are how Runx factors can accurately guide their contributions to distinct cell programs, whether by intrinsic DNA-binding sequence specificity, epigenetic constraints, or interactions with other partner factors. Here, we show how levels of expression of Runx itself control the qualitative choices of site occupancy to control T cell developmental speed and pathway choice.

Stages in T cell development are distinguished by changes in chromatin states and changes in expression of a discrete set of regulatory factors^20–26^, even though Runx factors themselves are collectively active at similar levels throughout^1^. Driven by Notch signaling and thymic microenvironmental cues, multipotent progenitor cells are converted to T-lineage committed pro-T cells within the thymus. They pass through CD4^-^CD8^-^ double negative (DN) substages (DN1-4) to CD4^+^CD8^+^ double-positive (DP) stage before becoming mature CD4 or CD8 single-positive (SP) T cells (Fig. 1a). Pro-T cells in DN1 and DN2a stages (“Phase 1”) still resemble hematopoietic stem and progenitor cells (HSPC) in regulatory gene expression and chromatin state and can still produce non-T lineage cells. Definitive T-lineage commitment normally occurs in transition from DN2a to DN2b stages. Up-regulation of T cell identity genes and changes in TF expression and chromatin states^20, 21, 23^ during commitment (DN2b) define “Phase 2”, extending till successful assembly of T cell receptor β (TCRβ) in DN3 stage (Fig. 1a). Runx1 and Runx3 are crucial for progression through both Phases. Note that although Runx1 and Runx3 act differently in other contexts^27^, they appear functionally redundant in pro-T cells^1^. However, they both bind to different genomic sites and regulate different target genes from Phase 1 to Phase 2^1^.

**Figure 1.**
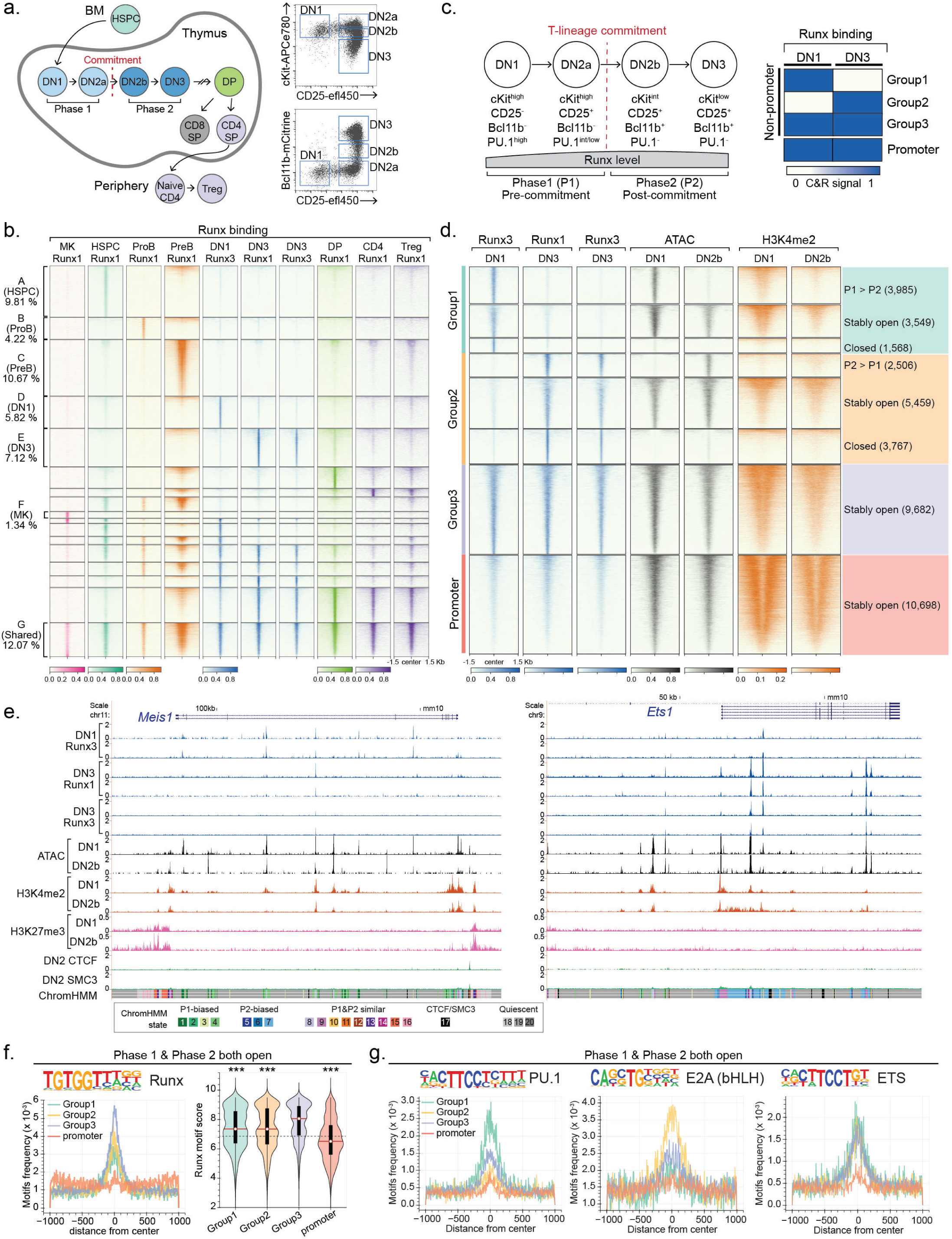
Runx TFs readily shift DNA binding site choice at different stages of T cell development largely independent of chromatin state changes. **a,** Schematic diagram shows different stages of T cell development. Hematopoietic stem and progenitor cells (HSPC), double negative (DN), double positive (DP), single positive (SP), and regulatory T cells (Treg). Representative flow cytometry plots (right) show cKit, CD25, and Bcl11b expression patterns in distinct DN populations. Note: “DN1” throughout refers only to cKit^high^ DN1, also known as Early T Progenitor (ETP) cells. Flow cytometry data was obtained from artificial thymic organoid (ATO) culture on day 9. **b,** Runx1 and Runx3 occupancy patterns in non-promoter regions of indicated cell populations are shown as peak-centered heatmaps. Runx binding profiles in DN1 and DN3 were detected by C&R from 2 independent experiments (merged data shown), and others were reported using ChIP-seq^15, 34–37^. MK, Maturing megakaryocytes. Numbers indicate the percent of group among total Runx binding sites. **c,** Diagram illustrates the key marker gene expression patterns in early T-cell development with associated levels of Runx1+Runx3 protein (left). Different groups of non-promoter Runx binding sites in early T-development are defined (right). **d,** Runx1 and Runx3 occupancy patterns in Phase 1 and Phase 2 cells are shown together with ATAC and H3K4me2 profiles^23, 24^. Stage-specific Runx binding groups were determined by C&R using DN1 (cKit^hi^ CD25^-^) cells obtained from *in vitro* OP9-Dll1 culture and thymic DN3 (cKit^low^ CD44^low^ CD25^+^) cells. **e,** Representative UCSC genome browser tracks for Runx C&R (independent replicates), and published ATAC-seq, and ChIP-seq data for H3K4me2, H3K27me3, CTCF, and SMC3 are shown^23, 24, 44^. Chromatin states computed by ChromHMM are displayed as a colormap at the bottom. P1, P2: Phase 1, Phase 2. Representative Phase 1-preferential Runx binding sites (near *Meis1*, left) and Phase 2-preferential sites (near *Ets1*, right) are displayed. **f, g,** Motif analysis was conducted within constantly open chromatin sites possessing different developmental patterns of Runx binding. **f,** Runx motif frequency within a peak (left) and the best Runx motif quality (position weight matrix score) within each peak (right) are shown. The dotted horizontal line on the violin plot indicates threshold motif quality to score as possessing Runx motif. Thin vertical black lines mark 1.5x interquartile range and thick vertical black lines show interquartile range. Red bars with white circles indicate median values. Two sample Kolmogorov-Smirnov (KS) tests, comparing each to Group3 motif scores: *** p<0.001. **g,** Motif frequencies of PU.1, E2A, and ETS factors in each Runx binding site Group are displayed.

Profound changes occur in chromatin looping, accessibility, and histone modification profiles during T-lineage commitment^22–24^, associated with repression of the genes associated with progenitor and alternative lineage states and activation of the T cell identity programs^20, 21, 28, 29^. Multiple TFs also change activity^21, 30, 31^. Runx TFs themselves can interact physically with multiple TFs at distinct binding sites, suggesting that TF cooperativity could be a major influence on Runx activity^13, 32, 33^. Yet how important are the Runx factors themselves in dictating which factor complexes and which target sites will be active?

Here, we evaluated the chromatin constraints on Runx action across the Phase 1 and Phase 2 stages of T cell development and tested the hypothesis that Runx binding site shifts depend on competition between Phase 1 and Phase 2 partners for a limiting amount of Runx protein. We found that at modest excess, when no longer titrated by Phase 1 partners, Runx factors directed a distinctive accelerated form of early T and innate-like cell development, relieving the need to repress most Phase 1 regulators before T-lineage regulatory genes could be upregulated. Both direct (binding site-mediated) and indirect mechanisms propagated through a Runx-dependent gene regulatory network drove this acceleration. Thus, Runx factor levels are major timing controllers of the deployment of the T-cell specification gene regulatory network.

## Results

### Runx TFs substantially shifted their binding sites during all stages of T-cell development

Runx 1 and Runx 3 are functionally redundant and bind to similar sites in early pro- T cells, but their site choices are highly stage dependent^1^. Fig. 1b shows that this not only distinguishes pro-T cell stages but also extends to Runx deployment in different hematopoietic lineage contexts from HSPCs to mature T cells, B lineage and megakaryocyte-precursor cells^15, 34–38^. Distinct genomic regions showed cell-type specific Runx occupancies, defining separate regions preferentially occupied only in HSPC, in B cell progenitors, in DN1, in DN3, in DP, or in mature T cells (Fig. 1b, A-F), and regions occupied in different developmental combinations, with ∼12% of sites shared in all (Fig. 1b, G). This site infidelity of Runx factors contrasted with binding patterns in pro-T cells of PU.1, a critical Phase 1-specific TF inherited from bone-marrow progenitor cells, which showed more similar binding site choices from HSPCs to DN2b pro-T cells (Fig. S1a). Importantly, each cluster of Runx binding regions from HSPC to mature T cells harbored a distinct set of motifs in addition to the Runx motif, in which motifs for EBF, PU.1, E2A, TCF1, GATA, JunB, and ETS factors were differentially enriched in each cluster (Fig. S1b). In our previous report, Runx binding was monitored by ChIP-seq after disuccinimidyl glutarate-assisted stabilization crosslinking^1^, which might have biased our previous results to overrepresent Runx complexes with other proteins rather than the distribution of Runx binding preferences themselves. Here, we independently analyzed Runx DNA binding profiles in pro T cells (DN1 and DN2b/DN3) using cross-linking-independent CUT&RUN instead^39, 40^ (C&R, Cleavage Under Targets & Release Using Nuclease)(see Methods; Fig. S1c-f shows detailed comparison of C&R against previous ChIP-seq identified sites). These results suggested that even excluding crosslinking artifacts, Runx factors readily changed their binding sites across all stages of T cell development to interact with distinct cell-type specific regions which may be occupied by different TF ensembles.

### Runx factors prefer “active” chromatin compartments but do not follow local chromatin state changes

We previously showed that some dynamic Runx binding shifts during T-lineage commitment occurred in a highly coordinated manner, with group appearance or disappearance of multiple Runx occupancies across large genomic domains of 10^2^-10^3^ kb (ref.^1^). To test whether Runx TFs were constrained or redirected by large-scale chromatin remodeling during commitment, the non-promoter Runx binding sites were categorized into three groups: Phase 1-preferential binding sites (Group 1), Phase 2-preferential binding sites (Group 2), and stably occupied sites (Group 3) (Fig. 1c). We analyzed how Runx binding was correlated with “active” (A) or “inactive” (B) large-scale nucleome compartments^41^ by comparing the principal component 1 (PC1) values of the previously reported Hi-C correlation matrices from ETP (DN1), DN2, and DN3 cells^22^. All Runx binding sites were preferentially enriched within the A compartment (84-92%) and were almost depleted from the B compartment (3-7%), regardless of Group (Fig. S2a, b; note *Ets1* flanking regions). As pro-T cells developed from ETP (DN1) to DN3 stages, most genomic regions remained in the same compartment (“A-to-A” 41.9%, “B-to-B” 48.1%). Among the minority of genomic regions changing compartment, Runx occupancy tended to follow the active states (Fig. S2a). Compartments switching from active to inactive (decreasing PC1 values, A to B trend) included more Group 1 (4.75%) than Group 2 sites (1.32%), whereas compartments becoming active (increasing PC1 values, B to A trend) included more Group 2 sites (4.56%) than Group 1 (2.35%) (Fig. S2a). One striking example of concerted Group 2 site appearance with a B to A compartment flip was seen in the extended flanking region of *Bcl11b* (Fig. S2b). However, most Runx site shifts occurred within A compartments.

Local chromatin states at numerous sites change substantially during the transition from Phase 1 to Phase 2, based on nucleosome density [assay of transposase accessible chromatin (ATAC), or DNase accessibility] and histone modifications^22–24^. To test how local chromatin state changes associate with Runx factor redistribution, we coded individual chromatin states across the genome from pre-commitment to post-commitment stages using ChromHMM^42, 43^ with published datasets for chromatin state marks in pro-T cells^23, 24, 44^(see Methods). Globally, Runx binding sites were not enriched in genomic regions with repression-associated chromatin states, whether defined by high levels of H3K27me3 or without any active chromatin marks (state 15, 16). Runx binding sites overall were preferentially enriched in open/active chromatin regions (state 1-3, 5-10) or weakly accessible regions harboring H3K4me2 marks representing likely poised regions (state 13)(Fig. S2c). This global bias was notable because we had verified that C&R could detect Runx binding in closed chromatin at least as well as ChIP-seq (Fig. S1f), and because Runx factors can work both as repressors and as activators^45–47^.

However, the developmental changes of Runx binding patterns did not strictly follow developmental changes in local chromatin states (Fig. S2c). At genomic sites occupied stage-specifically by Runx factors in Phase 1 or Phase 2 (Fig. 1d), developmental shifts in Runx occupancies could occur without corresponding changes in accessibility of those sites. Of Group 1 (Phase 1-specific) sites, only 43.8% were open in a Phase 1-restricted way, and only 21.4% of the Group 2 (Phase 2-specific) sites were open selectively in Phase 2. Therefore, over 50% of Group 1 and about 80% of Group 2 sites failed to change local chromatin accessibility as Runx binding changed in the Phase 1-Phase 2 transition. For instance, near *Meis1* multiple Runx occupancies disappeared from DN1 to DN3 (Group 1 peaks), but these sites remained open by ATAC-seq. Conversely, at *Ets1* multiple sites gained Runx occupancies from DN1 to DN3, but these sites had been accessible from DN1 (Fig. 1e, Fig. S2b). Over 1/3 of Group 2 sites remained closed in both Phases (Fig. 1d). Thus, local chromatin state failed to explain why Runx occupancy was delayed at Group 2 binding sites.

### Sequence specific features and partner factor interactions distinguish chromatin sites with developmentally changing Runx occupancies

Two other possible explanations for site choice shifts could be differences in Runx binding avidity (site affinity times site density) which could make Group 2 sites highly sensitive to small changes in Runx concentration, or collaborations with different stage-specific partners^1^. We evaluated these options by quantitative motif analysis, focusing exclusively on Runx sites that were consistently “open” to minimize chromatin effects (Fig. 1f, g). Runx binding sites mapping to open promoter regions had negligible Runx motif frequencies and poor Runx motif quality scores (Fig. 1f). Stably open non-promoter Runx binding regions in Groups 1, 2, and 3 had much higher Runx motif frequencies and motif quality than promoter sites, but to different degrees. Consistently occupied Group 3 sites had the highest scores. Both Group 1 and Group 2 sites showed lower Runx motif frequencies and motif scores than the Group 3 sites, but were similar to each other. Thus, at non-promoter sites without chromatin barriers, stage-specific redistribution of Runx factors occurred most readily between “modest” Runx motif sites without strong advantages for recruiting Runx factors via DNA recognition *per se* (Fig. 1f).

We previously identified distinct partner factors for Runx binding in Phase 1 and Phase 2^13, 32^, and found distinct partner motifs enriched at Runx sites in different stages^1^ (Fig. 1g, Fig. S1b). *De novo* motif enrichment analysis of the open sites confirmed that the Group 1 sites were highly enriched for PU.1 (ETS subfamily) motifs whereas the Group 2 sites were highly enriched for E2A (basic helix-loop-helix, bHLH) motifs. Although C&R Runx peaks did not show the extreme enrichment of ETS motifs seen with ChIP-seq (Fig. S1e), canonical (non-PU.1) ETS factor motifs were still enriched, and were found at similar frequencies in all classes of non-promoter sites (Fig. 1g). Thus, at sites that were stably accessible throughout Phases 1 and 2, different ensembles of TFs might recruit Runx TFs stage-specifically.

A central question is whether the T cell commitment process is switchlike, e.g. whether a single mechanism causes Runx factors to shift from Group 1 vs. Group 2 sites. The majority of precommitment-specific binding sites for Runx factors (Group 1 sites) have been shown to be actual co-binding sites with PU.1^1, 13^. Besides PU.1, in later pro-T cells other TFs such as GATA3 and Bcl11b can also collaborate with Runx factors at different sites^32, 33^. Notably, the presence of PU.1 can redirect Runx1 occupancy to the PU.1 sites, while depleting Runx1 binding (“theft”) from alternative Runx sites^13^. If Runx factor levels are truly limited such that partners have to compete to recruit Runx to different sites, the tipping of a balance between partners might cause concerted occupancy switches from Group 1 to Group 2 sites.

We hypothesized that if such competition occurs, it could be overridden if Runx availability were increased. We first tested this hypothesis in a PU.1 “theft” model (Fig. S3). The DN3-like Scid.adh.2C2 pro-T cell line, representing a Phase 2 state, was retrovirally transduced with exogenous PU.1, with or without additional exogenous Runx1 (Fig. S3a-b). PU.1 activated myeloid markers in the cells with or without exogenous Runx1 (Fig. S3c) and recruited endogenous Runx1 to a set of new co-occupancy sites with PU.1, most of which had been closed before (Fig. S3d, PU.1-induced). As previously reported^13^, without extra Runx1, PU.1 also caused a loss of Runx1 occupancy from nearly 55 % of the normal endogenous Runx binding sites (Fig. S3d, PU.1-depleted). However, when extra Runx1 was added (OE), although PU.1 was still able to recruit Runx binding to the PU.1-induced sites, occupancy of the PU.1-depleted sites was fully rescued (Fig. S3d). The extra Runx1 also occupied a set of novel sites (OE new). These had high quality Runx motifs at high frequency (Fig. S3e), but were mostly sequestered in closed chromatin in the normal Scid.adh.2C2. These results suggest that either Runx1-PU.1 complexes or high-level Runx1 alone could gain access to normally closed chromatin, but that the ability of PU.1 to remove Runx1 from its default binding sites was based on competitive titration when Runx1 was limiting.

### Modestly increased Runx1 in Phase 1 prematurely induced developmentally important TFs

If titration of potentially competing partner factors affects Runx site choice, Runx concentration might affect the T-lineage specification program in early progenitor cells. To test this, we exploited the OP9-Delta-like ligand 1 (Dll1) *in vitro* differentiation system^48^ as in our previous studies^1^. Briefly, bone-marrow derived progenitor cells expressing a *Bcl2* transgene and *Bcl11b*-mCitrine reporter were co-cultured with OP9-Dll1 cells and exogenous Runx1 was retrovirally delivered to pro-T cells when the progenitor cells were still at DN1 stage (Fig. 2a). Then, we measured T-development markers (cKit, CD44, and CD25) and *Bcl11b*-mCitrine expression, normally a marker for T-lineage commitment (see Fig. 1a, 2a)^49^. At day 2 after exogenous Runx1 introduction (overexpression, OE), total Runx1 protein levels were increased by 2-3 fold relative to the control conditions, and this increase was stable at day 4 post-infection (Fig. 2b, S3f). Notably, the modest degree of increase was important for the health of the cells^50^.

**Figure 2.**
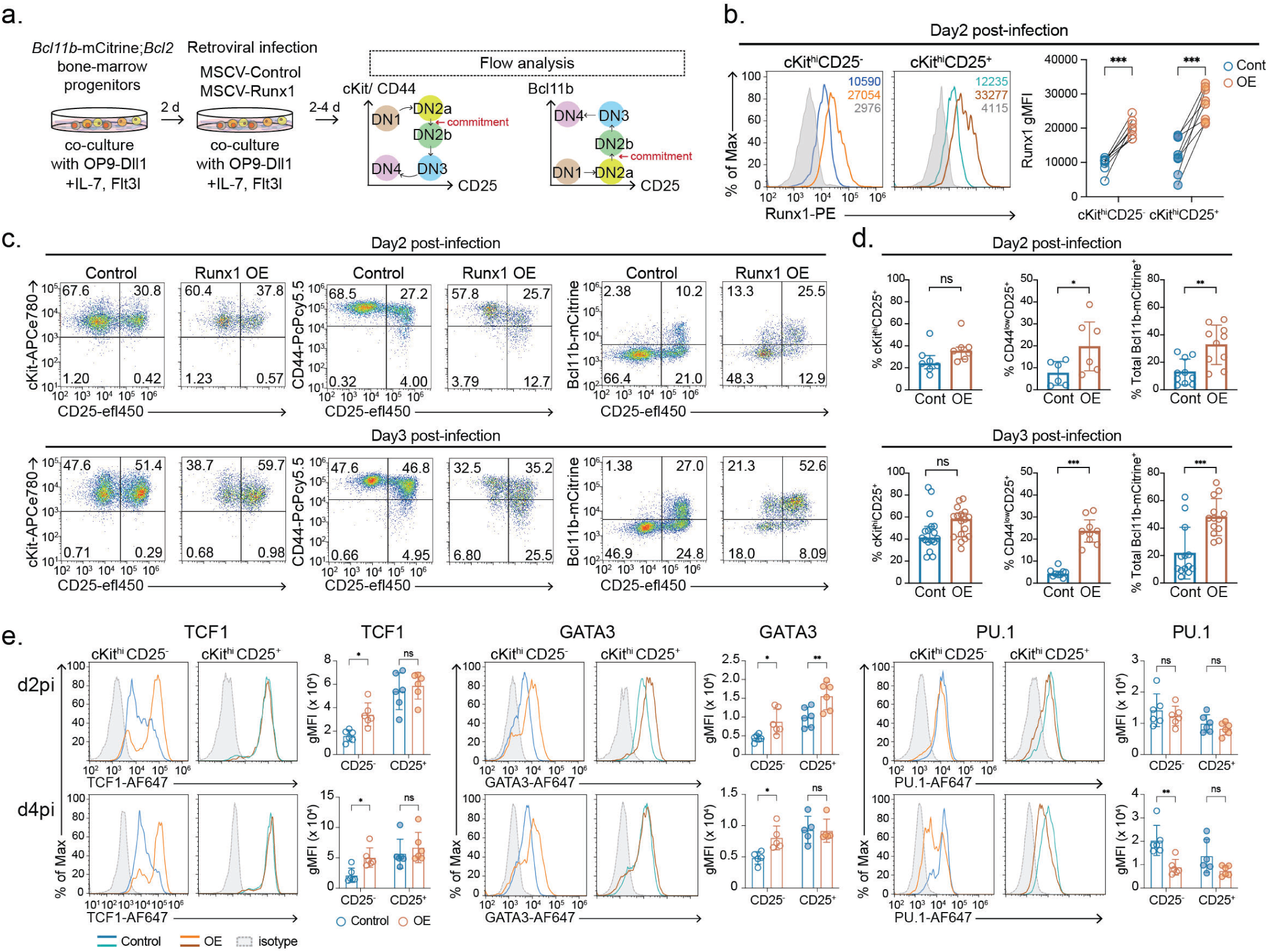
A mild increase in Runx1 level in Phase 1 prematurely upregulated Bcl11b, TCF1, and GATA3. **a,** Experimental design for testing Runx dosage effect on early T-development is displayed. **b,** Representative histograms show intracellular Runx1 protein levels detected by flow cytometry in control or Runx1-overexpression (OE) vector-transduced Phase 1 cells. Numbers indicate geometric mean fluorescence intensities (gMFI) of Runx1. Graph summarizes results from 8 independent experiments. Comparisons by two-way ANOVA. **c-d,** Flow cytometry data show cKit, CD44, CD25, Bcl11b-mCitrine reporter levels after delivering empty control or Runx1 overexpression (OE) vectors on day 2 (**c,** top) or day 3 (**c,** bottom) of T-cell development. Graphs in **d**, summarize mean values from 6-10 independent experiments with standard deviation (error bar). Comparisons by t-test. **e,** Histograms display protein expression levels of TCF1, GATA3, and PU.1 at day 2 and day 4 after Runx1 overexpression in Phase 1. Phase 1 (live, alternative lineage^-^, cKit^high^) cells were separated as CD25^-^ DN1 and CD25^+^ DN2 populations to compare target protein levels. Graphs display mean values from 5-7 independent experiments with standard deviations. Comparisons by two-way ANOVA. ***=p-value<0.001, **=p- value<0.01, *=p-value<0.05, ns=not significant.

Runx1 OE caused a striking acceleration of *Bcl11b* induction as early as day 2 after introducing extra Runx1, increasing at day 3: ∼20% of control cells but ∼50% of Runx1 OE DN2 cells showed *Bcl11b*-mCitrine expression (Fig. 2c, d). Abnormally, *Bcl11b*-mCitrine was also activated in some Runx1 OE DN1 cells (CD25^-^). Furthermore, increased Runx1 levels caused premature appearance of cells resembling DN3 cells (CD44^low^ CD25^+^)(Fig. 2c, d).

We examined expression profiles of developmentally important TFs, TCF1, GATA3, and PU.1. Runx1 OE upregulated TCF1 and GATA3 protein expression within the cKit^hi^ CD25^-^ DN1 (ETP) population at both days 2 and 4 post-infection (Fig. 2e). TCF1 and GATA3 levels in control cells only reached those of the Runx1 OE populations at day 4, in the cells that had turned on CD25 (DN2a cells)(Fig. 2e). PU.1 (*Spi1*) is normally repressed by Runx1 in Phase 2 only^1, 33^. Its expression was not affected by Runx1 overexpression at day 2 post-infection, but was significantly downregulated even in the DN1 population at day 4 (Fig. 2e), to levels lower than in normal CD25^+^ (DN2a) cells. Hence, a mild increase in Runx factor availability in Phase 1 pro-T cells could accelerate aspects of early T-cell development, especially within cKit^hi^ CD25^-^ DN1 cells.

### Runx level perturbations in Phase 1 resulted in profound changes in single-cell transcriptomes

We tested critically the ability of Runx factor levels to affect the T-cell specification program as a whole, using single-cell RNA-seq (scRNA-seq). To identify targets of Runx1 OE that were also dependent on normal Runx levels in controls, we also compared *Runx1/Runx3* double knockout (dKO) cells^1^, measuring the single-cell transcriptomes of Runx1 OE, control, and *Runx1/Runx3* dKO cells together using the 10X Chromium system. We delivered Runx1-OE vector or empty-vector control into *Bcl2*-transgenic progenitor cells to test OE, or guide-RNAs (gRNAs) against *Runx1* and *Runx3* or control irrelevant gRNAs into *Cas9;Bcl2* transgenic prethymic progenitor cells to test dKO. Then these progenitor cells were each co-cultured with OP9-Dll1 cells to two different timepoints (day 2 and day 4 post-infection for OE; day 3 and day 6 post-infection for dKO) and marked by hashtag oligos before they were pooled and subjected to single cell RNA-seq together (scRNA-seq) (Fig. 3a).

**Figure 3.**
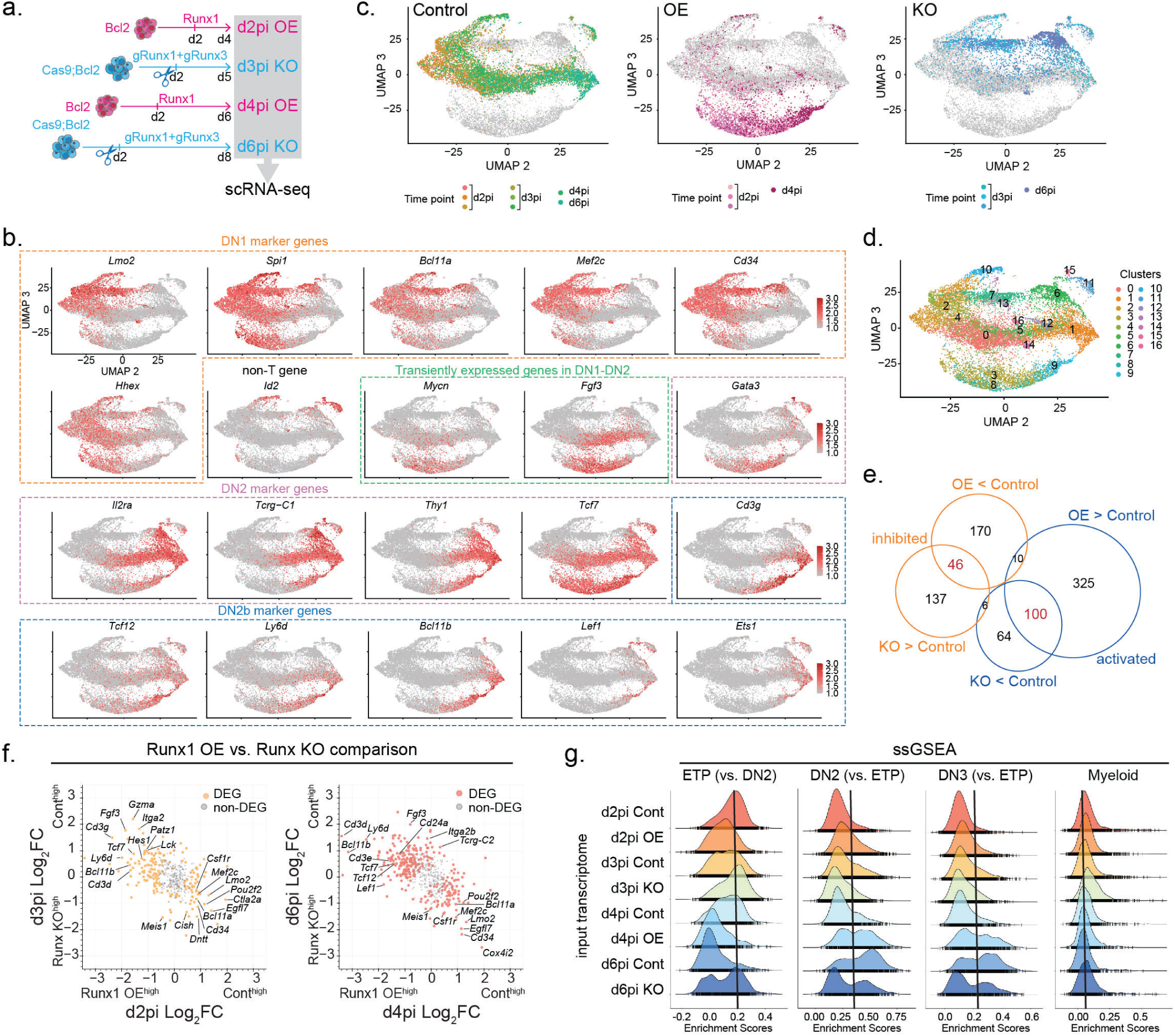
Single-cell transcriptomes revealed that Runx-level perturbation caused cells to take different developmental paths deviated from normal trajectory. **a,** Experimental schematics for single cell RNA-seq (scRNA-seq) are depicted. Each experimental condition group was marked by a different hashtag oligo (HTO) and equal numbers of cells from each were pooled for scRNA-seq. Two independent scRNA-seq experiments were performed. See Methods for details. **b-d,** UMAP2-3 illustrate scRNA-seq data from Runx1 OE and *Runx1/Runx3* double knockout (KO) in Phase 1. **b,** Color intensity in UMAP displays expression levels of indicated genes, which are informative to represent different T-development stages. **c,** Types of Runx perturbation (control, Runx1 OE, and Runx1/Runx3 KO cells) with cells from different experimental time points are highlighted in UMAP2-3 space. **d,** Cells from scRNA-seq are colored by Louvain clusters. **e,** Area-proportional Venn diagram shows the number of Runx-activated (blue) and - inhibited genes (orange) in OE and/or KO perturbations. The common, core target gene numbers are shown in red. **f,** Scatter plots compare Log_2_ fold-changes of Runx target gene expression in Runx1 OE and Runx KO conditions at d2-d3 (left) or d4-d6 (right) after Runx perturbations were introduced. **g,** Histograms display the aggregated enrichment scores of indicated pathways (ETP, DN2, DN3, and Myeloid pathways) in each cell computed from curated reference gene sets by ssGSEA. Cells were grouped by types of Runx perturbation and timepoints.

From two independent 10X runs, we recovered 15,310 cells that successfully passed the quality control criteria (see Methods). In a low dimensional transcriptome representation after batch correction, the first parameters in t-distributed stochastic neighbor embedding (tSNE) and Uniform Manifold Approximation and Projection (UMAP) reflected cell-cycle phases (Fig. S4a, b, top panels). After cell-cycle regression, UMAP2 (x axes, Figs. 3b-d) approximately represented real time developmental progression for normal pro-T cells, while UMAP3 (y axes) reflected perturbation; note that cells in each population progressed asynchronously. In controls, cells with low-UMAP2 values expressed high levels of DN1 signature genes (*Lmo2, Spi1, Bcl11a, Cd34, Mef2c, Hhex*). Genes transiently expressed during DN1 to DN2a transition (*Mycn, Fgf3*) were maximally expressed in control cells at UMAP2-intermediate values (Fig. 3b), while DN2 marker genes (*Il2ra, Tcrg-C1, Gata3, Tcf7, Thy1, Cd3g*) were first expressed in control cells at UMAP2-intermediate values and were maintained throughout the UMAP2-high cells. Then, mostly at later timepoints, genes associated with T-lineage commitment and the DN2b stage (*Bcl11b, Ly6d, Lef1, Ets1*) initiated expression in the UMAP2-high control cells (Fig. 3b). Thus, for controls, UMAP2 positions could relate cell states to the normal developmental progression.

### Developmental pathway kinetics were sensitive to the Runx dosage changes

Both Runx1 OE and *Runx1/Runx3* dKO (“Runx dKO”) caused contrasting deviations from the control cell clusters in the UMAP3 dimension (y axes in Figs. 3b-d). Control cells from all timepoints were concentrated at the center of UMAP3, whereas Runx1 OE cells were shifted to lower UMAP3 values while Runx dKO cells veered to UMAP3-higher values (Fig. 3c). Consistently, Runx perturbation caused cells to form unique clusters (Louvain clustering, Fig. 3d, S4c), suggesting that Runx factors regulated pathways followed by individual cells rather than changing subpopulation distributions along the control pathway.

Runx1 OE and Runx dKO cells showed evidence for reciprocally shifted landmark gene expression patterns along the UMAP2 axis as compared to the control group (Fig. 3b). Consistent with faster induction of *Bcl11b*-mCitrine reporter expression in absolute time, Runx1 OE cells upregulated *Bcl11b* at a lower UMAP2 value than control cells. In addition, Runx1 OE cells upregulated various later-stage genes “prematurely” at lower UMAP2 values and to higher levels than controls (*Gata3, Tcf7, Cd3g, Tcf12, Ly6d, Lef1*, *Ets1*). However, not all DN2-associated genes were concurrently induced (e.g., *Il2ra, Thy1, Tcrg-C1*), nor were all critical DN1 landmark genes downregulated (e.g., not *Spi1*) in Runx1 OE cells. On the other hand, Runx dKO caused lingering expression of DN1-associated genes (*Lmo2, Spi1, Bcl11a, Cd34, Mef2c*) with markedly impaired upregulation of later stage genes (*Mycn, Fgf3, Il2ra, Tcrg-C1, Gata3, Tcf7, Thy1, Cd3g, Bcl11b, Ly6d*). Instead, Runx dKO cells expressed genes associated with non-T cells, such as *Id2, Cd81*, *Csf2rb,* and *Ifngr2*, and prolonged expression of HSPC gene *Meis1* (Fig. 3b, Fig. S4d). Thus, many developmentally regulated genes sensitively responded to perturbations of Runx levels, although alternative programs were not coherently activated or inhibited together. These results also indicated that Runx dosage responses occurred within pro-T cells themselves, not only reflecting enrichment of minority contaminants.

### The common set of genes sensitive to both gain and loss of Runx functions overlapped with essential genes for early T cell development

We reasoned that the core set of development-regulating genes that were most likely to be direct Runx targets should be reciprocally affected by Runx1 OE and *Runx1/Runx3* dKO. Differentially expressed genes (DEGs) affected by Runx1 OE and *Runx1/ Runx3* dKO were each similar at both perturbation timepoints (Fig. S4e). The global gene expression changes mediated by Runx dKO vs. Runx1 OE were inversely correlated at both timepoints, with a core set making significant, reciprocal responses to both gain- and loss-of Runx functions (Fig.3e, f and S4f, Table S1, S2). These core Runx- activated genes (100 genes) were generally upregulated as normal pro-T cells advance to the DN2 and DN3 stages (e.g., *Cd24a, Hes1, Patz1, Ahr, Myb, Lck, Tcf7, Ly6d, Bcl11b, Lat, Cd3d, Cd3g,* and *Gzma*). In contrast, core genes inhibited by Runx factors (46 genes) included ETP signature and non-T genes (e.g., *Mef2c, Lmo2, Cd34, Pou2f2, Csf1r, Csf2rb, Bcl11a, Id1, Ly6a,* and *Cd81*)(Fig. 3e, f, and S4f).

These impressions from landmark genes were supported globally by Single-sample Gene Set Enrichment Analysis (ssGSEA)(Fig. 3g). GSEA used curated stage-specific thymocyte gene sets to track developmental progression at different absolute times of differentiation, shown in time-resolved population histograms. Controls started with high ETP (DN1) and low DN2 or DN3 enrichment scores and shifted to low ETP (DN1) and high DN2 and DN3 values from day 2 to day 6 post perturbations. Runx dKO cells, at both day 3 and day 6, were more significantly correlated with ETP signatures and slightly increased myeloid signatures, failing to activate DN2 or DN3 signatures comparably to controls. In contrast, Runx1 OE at both timepoints showed accelerated loss of associations with ETP and increases in DN2, DN3 profiles accelerated by about a day relative to controls (Fig. 3g).

Thus, single-cell transcriptome analysis suggested that Runx activity preferentially activated genes critical for T-developmental progression, while inhibiting the genes associated with progenitor and myeloid programs.

### Runx levels controlled T-developmental speed via selective activities on discrete T-identity and lymphoid program modules

The fine structure of Runx effects on developmental progression could be measured quantitatively using pseudotime. We calculated pseudotime trajectories with Monocle3, defining the root cells by high expression of *Flt3* and *Kit* and absence of *Tcf7* and *Il2ra* transcripts, based on the phenotype of the earliest thymus-seeding ETPs^25, 51, 52^. As expected, from day 2 pi to day 6 pi, cells in control clusters showed gradual pseudotime progression with UMAP2 value (Fig. 4a, b). Runx dKO displayed slower progression in pseudotime as compared with control cells, while Runx1 OE markedly accelerated progression (Fig. 4a. b).

**Figure 4.**
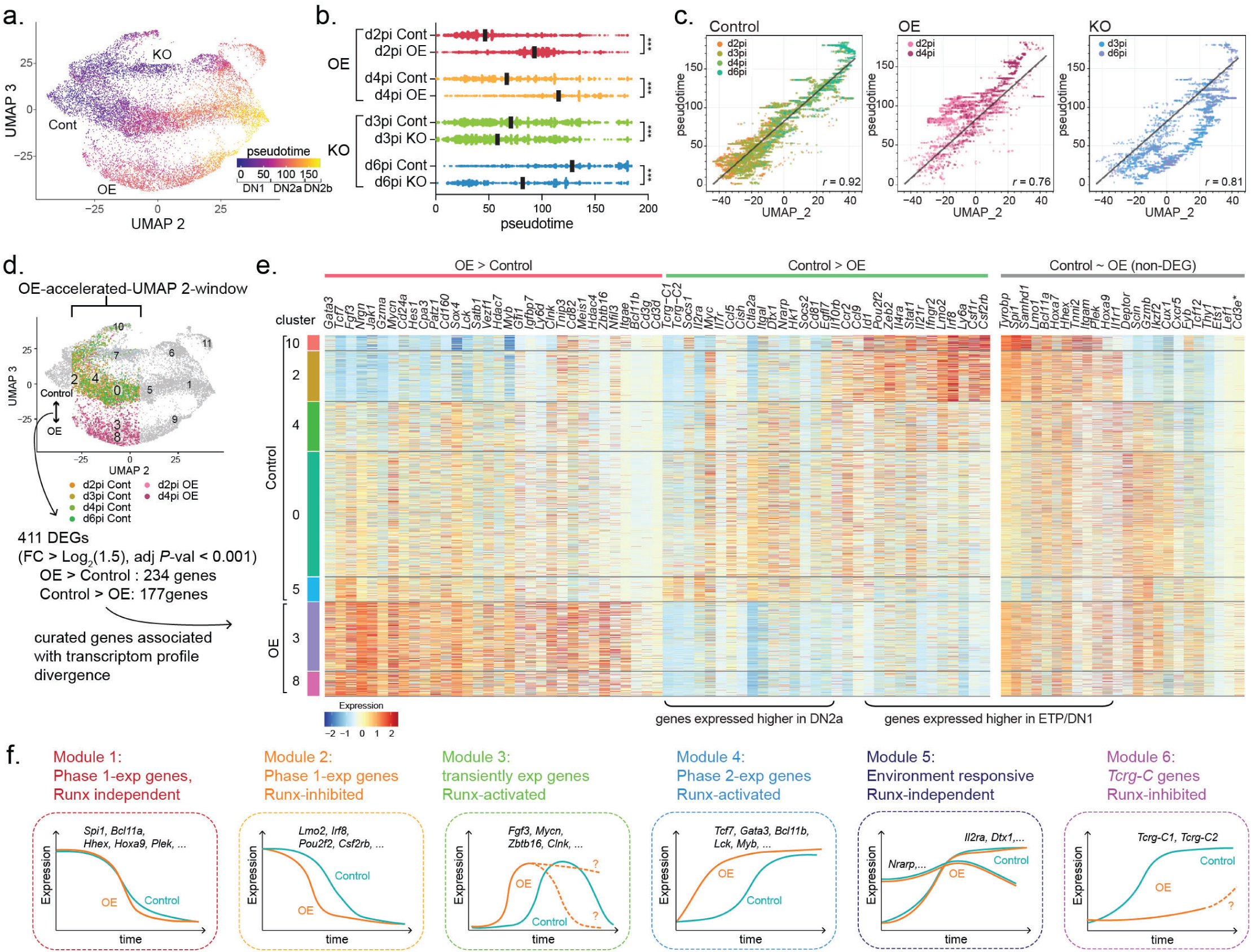
Runx levels control T-development progression rate by activating selective gene network modules. **a,** The pseudotime score of each cell is displayed in UMAP2-3 by color. Pseudotime score was calculated with Monocle 3 by defining the principal root node with cells expressing high levels of *Flt3* and *Kit* and absence of *Il2ra* and *Tcf7* transcripts. **b,** Pseudotime distributions of cells from indicated groups are shown with median using a black bar. Kruskal-Wallis test of multiple comparisons. ***adj.p-value<0.001. **c,** Scatter plots compare the pseudotime score and the UMAP2 value, which approximately corresponds to the real time. Cells in control (left), OE (middle), and KO (right) groups are shown with Pearson correlation *r*. Black line indicates a linear regression fit calculated on control group. Spearman’s rank correlation coefficient ρ was also computed: control (ρ = 0.906, p<0.0001), OE (ρ = 0.733, p<0.0001), KO (ρ = 0.819, p<0.0001). **d,** Analysis strategy for differential gene expression is shown. **e,** Curated list of differentially expressed genes between control vs. Runx1 OE groups within −30 < UMAP2 < 5 window is displayed in heatmap (left). Genes that were developmentally dynamically regulated (defined by cluster 1 vs. cluster 2 comparison), yet not differentially expressed by Runx1 OE within −30 < UMAP2 < 5 cells, are also shown (right). \**Cd3e* was scored as a non-DEG due to low frequency of Runx1 OE cells expressing *Cd3e* at this early timepoint. **f,** Graphical illustration of gene expression modules utilized in early T-cell development.

For control cells, UMAP2 and pseudotime parameters were correlated, with a highly linear relationship (Pearson correlation score *r* = 0.92) (Fig. 4c), but Runx dKO and Runx1 OE samples showed weaker linearity than the controls (Fig. 4c middle and the right panels). Runx dKO cells exhibited consistently slowed developmental (pseudotime) progression across most UMAP2 values; in contrast, Runx1 OE cells accelerated development (pseudotime), especially during a specific low-UMAP2 window (“OE- accelerated-UMAP2-window”: UMAP2: −30 to 5). This implied a specific early-Phase 1 window of opportunity when elevated Runx1 dosage was most effective.

To define the target genes involved, we compared differentially expressed genes (DEGs) between control vs. Runx1 OE groups specifically within the stages when they presumably diverge, focusing on cells within the same “OE-accelerated-UMAP2-window” (−30 < UMAP2 < 5) (Fig. 4d), approximately corresponding to clusters 4, 2, and 0 (control) vs. clusters 3 and 8 (Runx1 OE). Among 411 DEGs, 234 genes were more highly expressed in the OE group than the control, while 177 were higher in the control group (Table S3). The genes upregulated by Runx1 OE in this focused comparison included multiple key T-identity genes and driver TFs (*Cd3g, Cd3d, Bcl11b, Tcf7, Lck, Gata3, Myb, Patz1, Hes1*), which were induced not only much earlier but also at higher levels than in controls. However, Runx activation targets were not entirely T-lineage specific, as Runx1 OE also caused increased expression of genes (*Zbtb16, Nfil3, Clnk, Cd160*) associated with innate lymphoid cell (ILC) programs, in which Runx3 is more highly expressed^53^. These genes are normally transiently activated in DN1-DN2a cells but repressed during later T cell development by Bcl11b and E-proteins (Fig. 4e, Table S5)^32, 54, 55^.

The genes more highly expressed in control than in Runx1 OE samples were also striking, as they were enriched for cytokine-associated and Notch signaling responsive genes (*Il2ra, Il7r, Il4ra, Il21r, Stat1, Socs1, Socs2, Cish, Dtx1, Nrarp, Myc*) and for TCRγ- constant region genes (*Tcrg-C1, Tcrg-C2*), which are also positively regulated by cytokine signaling^56, 57^. Of major Notch target genes, only *Hes1* was upregulated by Runx1 OE. Whereas these other genes normally increase expression during ETP to DN2b progression, it was notable that increased Runx1 availability activated T-identity and common lymphoid program genes without inducing these environmental signaling response genes. Finally, some Phase 1 genes (*Lmo2, Irf8, Pou2f2*) were prematurely inhibited by Runx1 OE. However, other Phase 1 genes including key TFs, *Spi1, Meis1*, *Hoxa9*, *Hhex,* and *Bcl11a* were not prematurely turned off within this pseudotime window.

This uncoupled, selective target gene activation by Runx1 OE was intriguing, as the Runx target genes included TFs which are known to be potent at directing early T-cell development. Together, these results indicated a modular structure of pro-T cell gene regulatory network topology in which distinct subprograms were not necessarily coherently linked (Fig. 4f). Runx TFs exerted selective activities on inducing T-identity, shared, and ILC-specific programs without activating cytokine/proliferation programs nor completely blocking the stem and progenitor program, yet still accelerating T-developmental progression from DN1 to DN2-like stages.

### Runx1 overexpression supported NK cell potential while inhibiting myeloid and granulocyte programs in the absence of Notch signaling

As Runx factors appeared to activate genes associated not only with T cell identity, but also with ILC lineage, we asked whether increased Runx availability in pro-T cells changed developmental potentials, assayable in the absence of Notch signaling. To evaluate alternative lineage potentials, we designed a competitive commitment assay by introducing control empty vector or Runx1-OE vector, which were distinctively marked by mCherry or human NGFR (hNGFR) expression. Then, we sorted the same numbers (100 cells per well) of transduced DN1, Bcl11b^-^ DN2a, and Bcl11b^+^ DN2a cells each from control- and Runx1-OE populations and co-cultured them with OP9 stroma, either expressing Dll1 to assess T cell potential or without Dll1 (OP9-Control) for alternative potential (Fig. S5a). Fig. S5b shows live cell frequency and number for each condition after 6 days. Runx1 OE cells yielded lower overall cell recoveries and frequencies relative to controls for both Notch-dependent and -independent conditions. The disadvantages in cellular proliferation and/or survival of Runx1-OE cells could have reflected their impaired activation of cytokine/signaling-pathways and lower expression of *Myc* (Fig. 4e, g).

Elevated Runx levels caused qualitative differences in alternative lineage choices under Notch-independent conditions (Fig. S5c). In DN1 stage, the granulocyte/myeloid lineage path is a common alternative for control cells, but Runx1 OE disfavored this. Instead, Runx1 OE DN1 cells preferentially diverged to express NK cell marker (NK1.1). Unexpectedly, Bcl11b^+^DN2 cells from Runx1 OE samples could still upregulate NK1.1 in the absence of Notch signaling, which was blocked in control Bcl11b^+^ DN2 cells (Fig. S5c). This could be ascribed to the precocious onset of *Bcl11b* expression, potentially before commitment could be complete, along with increased levels of ILC-associated genes in Runx1 OE cells. Together, our data show that moderately raised Runx levels supported NK cell-associated programs, while counteracting myeloid/granulocyte potentials.

### Modestly increased Runx1 drove faster T-developmental progression until DN4

The single cell transcriptome profile suggested that Runx levels had a significant impact on early T-developmental progression on the pseudotime trajectory. However, effects on cytokine and Notch-signaling response programs and on key TFs supporting Phase 1 were not coordinated with these changes as they are in normal cells, which could also promote deviation from the normal developmental pathway. Furthermore, as Runx1- OE DN2 cells expressing Bcl11b still were incompletely committed, unlike normal Bcl11b^+^ DN2 cells (Fig. S5c), the question remained whether Runx1 overexpression truly drove faster T cell development. To track long-term developmental consequences, we took advantage of a three-dimensional (3D) artificial thymic organoid (ATO) system using mouse MS4-Dll4 feeder cells, which closely recapitulates thymic T-cell developmental stages from DN1 to CD4- or CD8- single-positive stages^58, 59^. We formed mixed chimeric ATOs, mixing the same numbers (1,000 input cells) of bone marrow progenitor cells transduced with control- or Runx1 OE-vectors marked with either mCherry or hNGFR, as shown in Fig. 5a, and we then compared their T-developmental progression within the same ATOs on day 5, 8, 10, and 15.

**Figure 5.**
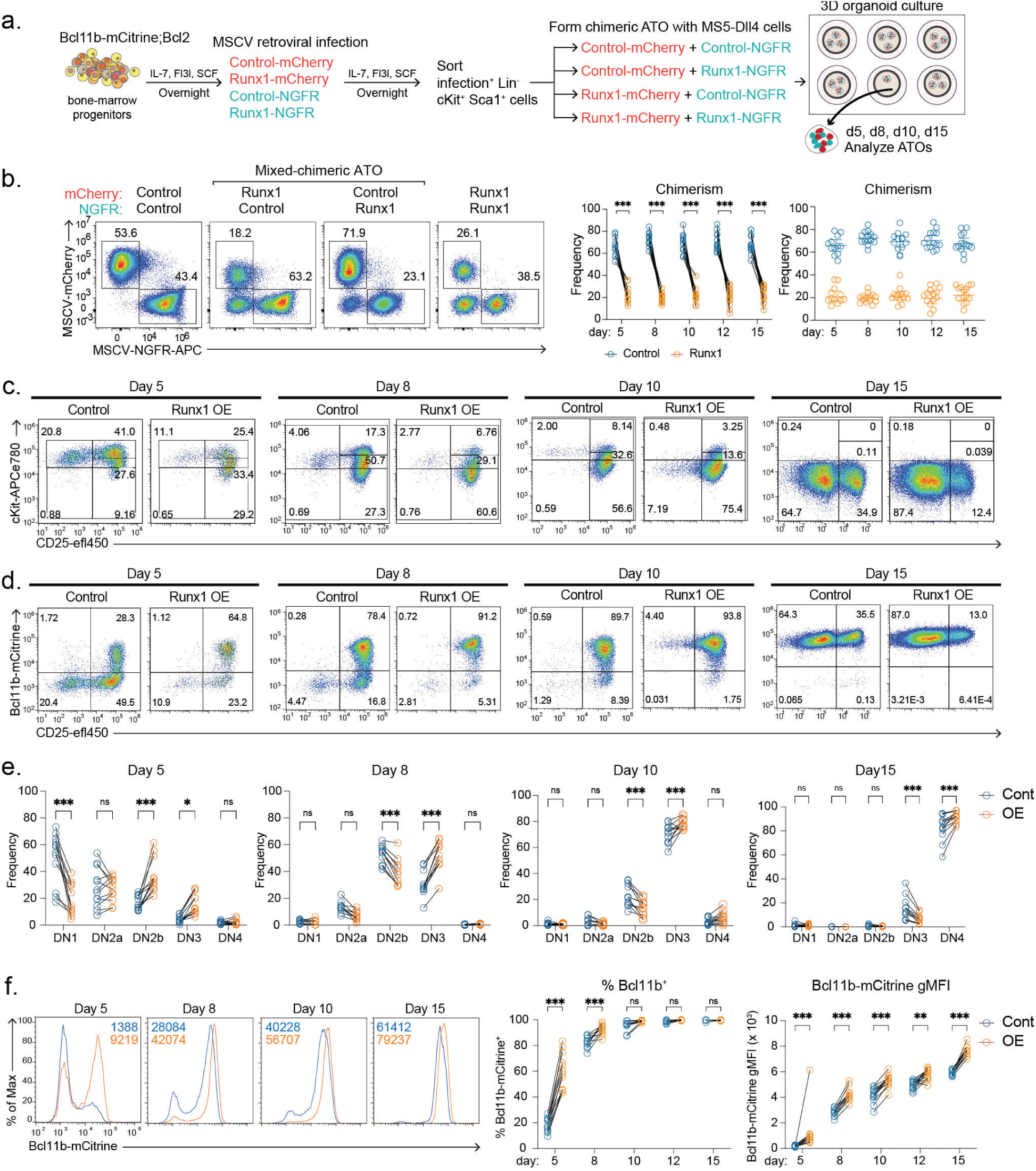
Runx1 overexpression results in overall faster T-lineage development from DN1 to DN4 stages in the mixed chimeric artificial thymic organoid. **a,** Experimental schematics for mixed-chimeric artificial thymic organoid (ATO) culture are illustrated. **b,** Representative flow plots display expression levels of infection markers (gated on live lineage^-^ CD45^+^ cells). Graphs show average frequency of control vs. Runx1 OE cells from chimeric ATOs at indicated timepoints. Comparisons by two-way ANOVA. **c-d,** Expression profiles of T-development markers, cKit, CD25, Bcl11b-mCitrine were measured by flow cytometry. Representative plots were gated on indicated infection marker^+^ cells at indicated timepoints. **e,** Graphs show frequencies of indicated pro-T cell populations in control- or Runx1-OE transduced cells at different time points. 3 independent experiments, n=11-14 ATOs. **f,** Bcl11b-mCitrine reporter expression levels during d5-day15 ATO cultures were shown. Numbers in histograms (left) indicate gMFI. Graphs show percent of Bcl11b-mCitrine+ cells (middle) and Bcl11b-mCitrine gMFI (right). Two-way ANOVA: ***=p-value<0.001, **=p-value<0.01, *=p-value<0.05, ns=not significant.

As under the conditions from the competitive commitment assay, Runx1 OE progenitor cells showed only ∼20% of chimerism at day 5 post-culture, whereas control progenitor cells comprised at least ∼60-70% of the populations (Fig. 5b). However, the frequencies of Runx1 OE progenitor cells did not decrease further at the later timepoints, and a similar chimerism was stably maintained until the end of the analysis (day 15).

Although Runx1 OE resulted in lower cell recovery than the control group, we observed a striking T-developmental acceleration. At day 5, about 65% of cells turned on *Bcl11b* in the Runx1 OE condition, progressing to DN2b and DN3 stages, and only ∼10% cells remained at DN1 stage (Fig. 5c-e). In contrast, ∼20% and ∼50% of control cells were still at the DN1 and DN2a stages respectively. This faster development by Runx1 OE continued through later stages, as they progressed to DN3 stage faster (on day 8 and day 10), and reached DN4 stage earlier (on day 15) than the control group (Fig. 5c-e). Moreover, Runx1 OE not only advanced Bcl11b onset, but also increased Bcl11b expression per cell at all timepoints even beyond DN3 stage (Fig. 5f), extending previous evidence^49^. Thus, increased Runx1 levels in progenitor cells propelled intrinsically faster T-cell development to DN4 stage, with prominent acceleration especially across the DN1 to DN2b transition.

### A modest increase in Runx1 protein levels in Phase 1 resulted in premature Runx1 occupancy in post-commitment-specific sites

To understand how elevated Runx1 levels drove faster T-developmental progression, Runx1 binding profiles were examined in Phase 1 pro-T cells sorted (Lineage^-^, infection^+^, CD45^+^ cKit^high^ cells) 40-42 hr after Runx1 or control vector introduction (Fig. S6a). C&R analysis showed a clear increase in the number and intensities of Runx1 occupancies across the genome in Runx1 OE cells as compared to control cells transduced with empty vector (Fig. 6a, Fig. S6b, c). As these cells were stillmostly in Phase 1 (cKit^high^) at harvest, Runx1 occupancies in the empty vector control group (Fig. 6a, Fig. S6c, Cont Runx1) were similar to Runx3 binding in unperturbed DN1 cells (Fig. 6a, DN1 Runx3). Such Group 1 and Group 3 sites were also strongly occupied in the Runx1 OE cells (Fig. 6a, S6c, d top row). However, the OE samples also showed new Runx1 occupancies at two classes of non-promoter sites. A notable subset overlapped with 65% of normal Group 2 sites (Group 2a, Fig. 6a, Fig. S6c). Examples of Group 2a sites precociously occupied when Runx levels were elevated were found in the *Bcl11b* enhancer region, as well as in Runx-responsive Runx loci *Ets1*, *Cd3* cluster, *Tcf7*, *Thy1,* and *Zbtb16* (Fig. 6b, S6d bottom row). These Group 2a sites were thus conditionally accessible in Phase 1, depending on Runx1 availability, distinct from the remaining Group 2 sites (Group 2b, Fig. 6a).

**Figure 6.**
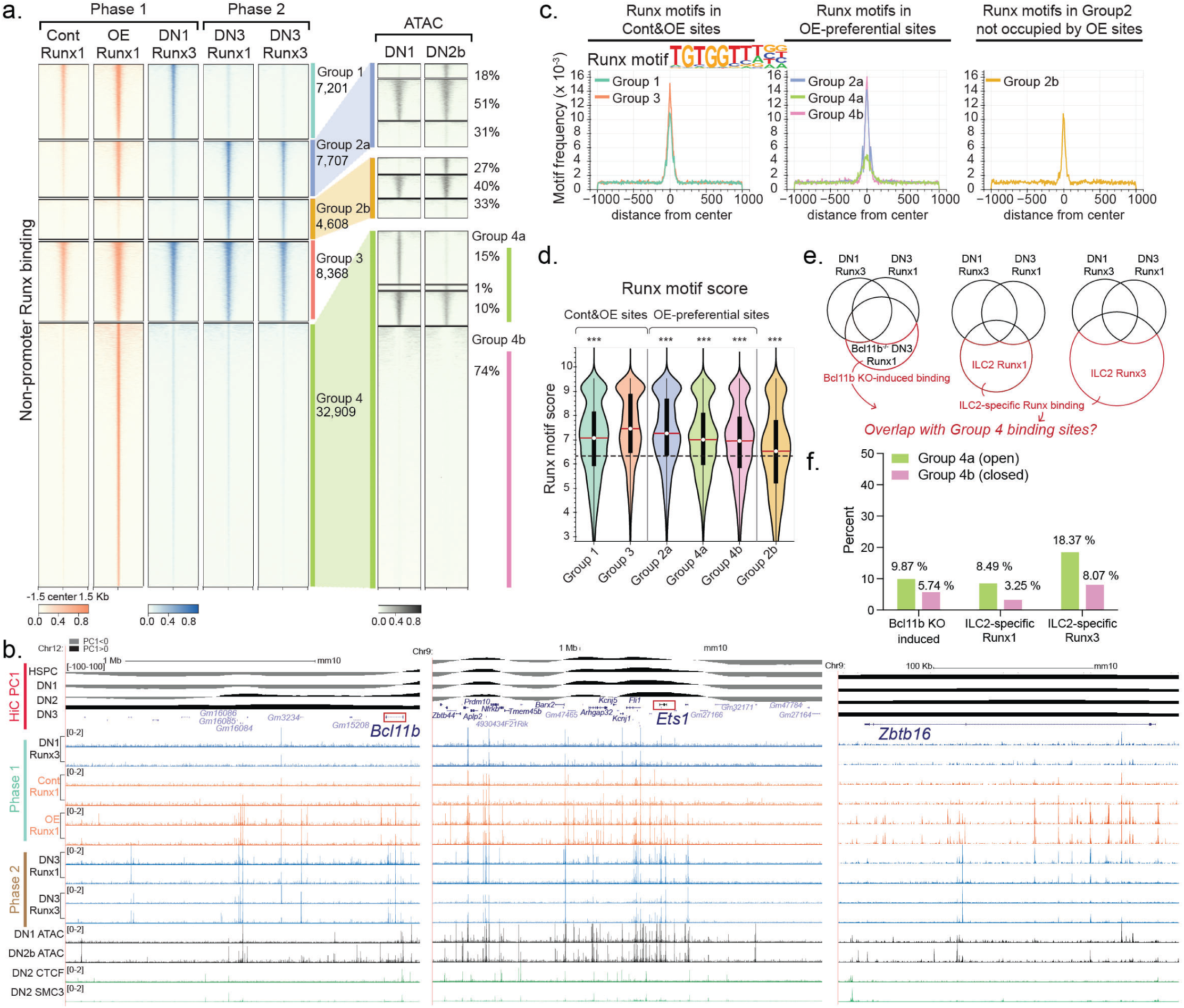
A modest increase of Runx1 concentration resulted in premature occupancy in post-commitment-preferred sites and new sites. **a,** Heatmap represents Runx1 or Runx3 DNA binding patterns in non-promoter regions from indicated cells. Orange tracks were derived from experimental cells and blue tracks were obtained from unperturbed Phase 1 (*in vitro* DN1) and Phase 2 (thymic DN3) pro-T cells (two independent C&R experiments for each condition). Stage-dependent chromatin accessibility patterns in normal cells^23^ at Group 2a, Group 2b, and Group 4 sites are shown on the right with percent of total peaks in a group. **b,** Representative UCSC genome browser tracks display Runx binding (C&R) together with published HiC PC1 values, chromatin accessibility (ATAC) profiles, and binding sites of loop forming machinery (CTCF and SMC). Enhancer regions near *Bcl11b*, *Ets1*, and *Zbtb16* are displayed. **c,** Runx motif frequencies in different Groups of Runx binding sites are illustrated as density plots. **d,** Violin plot demonstrates the best Runx motif score distribution in each Groups of Runx binding sites. Two sample KS test (comparing to Group3 motif scores). *** p<0.001. The horizontal dotted line shows the threshold PWM score to be considered to harbor the Runx motif. Thin vertical black lines mark 1.5x interquartile range and thick vertical black lines show interquartile range. The red lines with white circles indicate median values. **e,** Testing hypothesis that Runx1 OE accesses sites conditionally occupied in other pro-T related contexts. Area-proportional Venn diagrams show analysis strategy to identify Runx binding sites appearing specifically in Bcl11b knockout DN2b/DN3 cells (left), and ILC2-specific Runx binding sites (middle; Runx1, right; Runx3). **f,** Bar graph shows percentages of Group 4 peaks overlapping with indicated Runx binding site types.

Most Group 2a sites, like Group 2 sites generally, showed unchanging ATAC- profiles in normal development (51% constantly open, 31% stably closed in Phase 1 and Phase 2); only 18% of these sites gained accessibility after T-lineage commitment. However, Group 2a sites showed different Runx motif qualities from Group 2b, as Group 2b sites had lower-quality and less abundant Runx motifs compared to Group 2a sites (Fig. 6c, d). Thus, Runx level itself was insufficient to accelerate binding to Group 2b sites.

Overexpressed Runx1 also bound sites that were normally unoccupied in primary pro-T cells (Group 4, Fig. 6a), as expected (cf. Fig. S3). In contrast to other Runx binding sites, Group 4 sites were largely inaccessible normally (75%, Group 4b); only 25% were open in Phase 1 (Group 4a)(Fig.6a). Here, Group 4a and 4b sites had lower motif quality than the Group 2a sites (Fig. 6d), but Group 4b sites had a higher Runx motif density than Group 4a sites (Fig. 6c), suggesting that closed chromatin requires more numerous Runx motifs than open chromatin to engage Runx factors. However, at larger scales, all groups of Runx binding sites including Group 4 regions were still mainly associated with A compartments, suggesting that increased Runx levels could not overcome the inactive compartment barrier (Fig. S6e).

As Group 4 sites were not normally occupied in normal pro-T cells, we asked if they overlapped with Runx binding sites appearing only 1) in Bcl11b-KO DN3 cells^32^ (Bcl11b KO-induced sites) or 2) in ILC2 cells^53^. Indeed, ∼18% of Group 4a sites corresponded with Runx3 binding sites specific to ILC2, suggesting that *de novo* OE-specific Runx binding sites in open chromatin included some ILC-associated regulatory regions.

### Stage-specific and dosage-dependent Runx binding patterns near the Runx target genes

As Runx target genes encoded many developmentally important TFs, Runx dosage-sensitive effects might reflect cooperation with other TFs. Such cooperating factors might either bind directly with Runx factors to guide them to functionally important sites, or could work independently on a separate set of targets. To distinguish these modes of action, we evaluated Runx-OE accessed sites for their potential association with Runx-activated factors.

The most frequently detected motifs showed distinct enrichment profiles in each site Group (Fig. 7a). PU.1 motifs were not only enriched in all Phase 1-occupied binding sites as expected (Group 1 and Group 3), but also in the open Group 4 binding sites (Group 4a), though not in any Group 2 sites. Conversely, TCF1/HMG motifs were frequent in both types of OE-preferential sites (Group 2a and Group 4), but sparsely discovered in Group 1 sites. E2A (bHLH) motifs were only enriched in Group 2 sites, but interestingly, most highly in those that were not OE-accessible (Group 2b). Finally, GATA motifs were generally not enriched in Runx binding sites (Fig. 7a). These results suggest that stage-specific and/or dosage-dependent Runx binding sites may be co-occupied with different pools of TFs.

**Figure 7.**
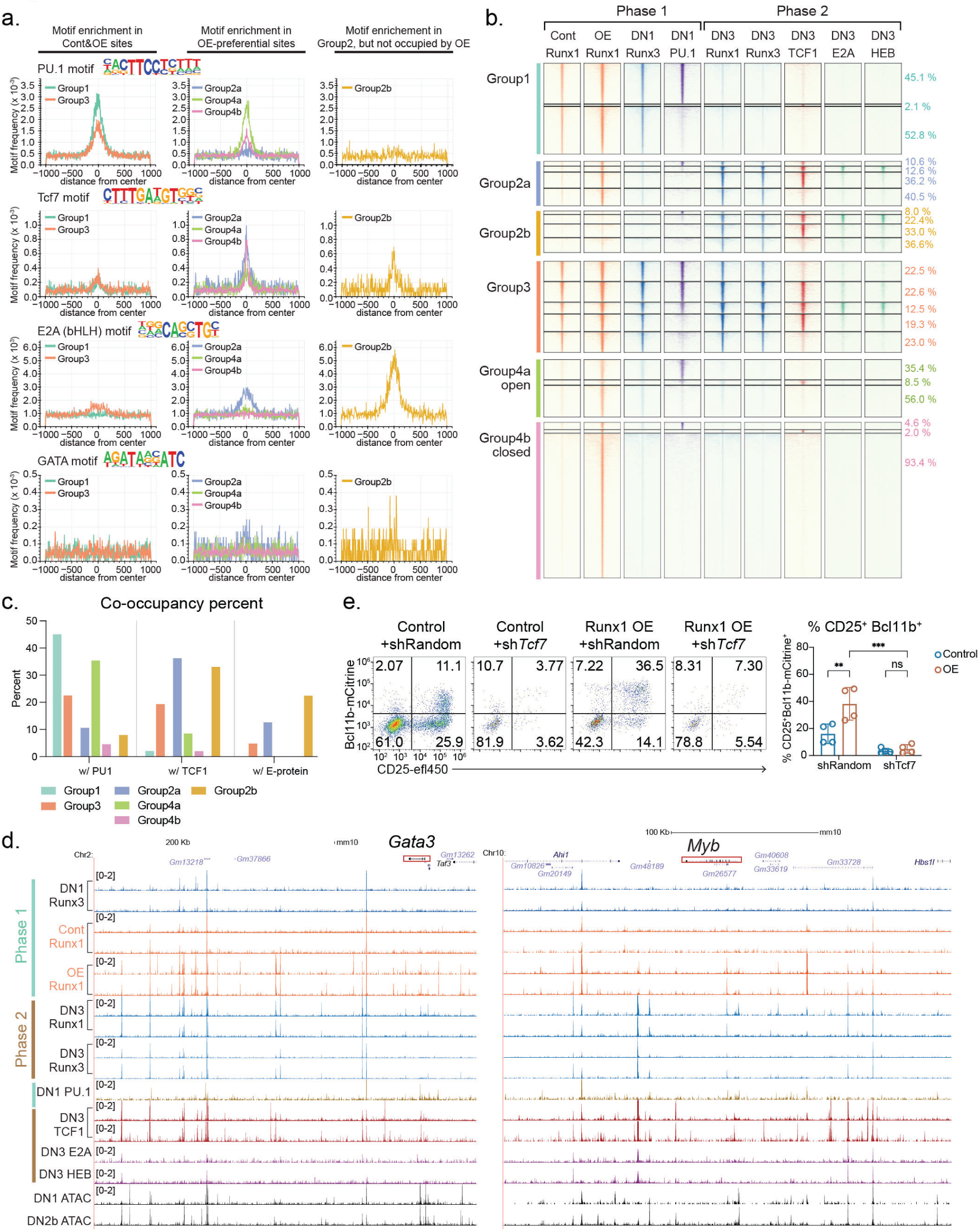
Runx factors engage functional target gene regions together with PU.1, TCF1, and E-proteins. **a,** Density plots illustrate motif frequencies for PU.1, TCF1 (Tcf7), bHLH, and GATA factors in different types of Runx binding sites. **b,** Runx1, Runx3 (blue), PU1 (purple)^24^, TCF1 (red), E2A and HEB (green) binding profiles in non-promoter regions under unperturbed Phase 1 or Phase 2 conditions are shown. Runx1 binding patterns in empty vector control and Runx1 OE transduced conditions are displayed in orange tracks (left). Stage-preferential dynamic binding groups are indicated as color bars. Group 1, Phase 1-preferential; Group 2a, Phase 2-preferential and precociously occupied by OE; Group 2b, Phase 2-preferential but not occupied by OE; Group 3, Phase 1 & Phase 2 shared; Group 4a, OE-specific and open sites; Group 4b, OE-specific and closed sites. The numbers on the right side indicate percent of each group of peaks within the same color bar. TCF1, E2A, and HEB binding sites were measured in independent replicates using C&R from thymic DN3 cells. PU.1 occupancy was previously determined using ChIP-seq^24^. **c,** Number of Runx binding sites co-occupied with PU.1 or TCF1 or E-proteins were enumerated and their percentages in each group are shown using a bar graph. **d,** Representative UCSC genome browser tracks near *Gata3* and *Myb* show indicated TF binding profiles. **e,** shRNA against *Tcf7* or random control shRNA was introduced to bone-marrow progenitor cells in combination with Runx1-OE or empty control vector, then the progenitor cells were co-cultured with OP9-Dll1 for 2 days. Bar graph summarizes Bcl11b-mCitrine and CD25 expression levels measured by flow cytometry with mean and standard deviation. n=4 independent experiments, Two-way ANOVA. ***=p-value<0.001, **=p-value<0.01, ns=not significant.

Actual binding of partners, assessed by C&R in their peak stages of action, showed more substructure than these motif enrichment-predicted patterns. PU.1 occupied ∼45% of Group 1 sites in Phase 1 cells, while TCF1 occupancy (measured in Phase 2 cells) was rarely (∼5%) detected in Group 1 sites. Group 2a and Group 2b sites did not overlap with PU.1 binding sites, but >50% of both Group 2a and 2b sites were co-occupied with TCF1 in Phase 2 cells. Distinct from PU.1 and TCF1, two E-proteins highly expressed in pro-T cells, E2A and HEB, were detected binding only at a small fraction of Group 2 sites, in each case sharing occupancy with TCF1, as shown by examples of enhancer regions of *Gata3* and *Myb* (Fig. 7d). Therefore, PU.1 in Phase 1 and TCF1 and E-proteins in Phase 2 interacted with specific subsets of Group 1 and Group 2 Runx binding sites, respectively (Fig. 7b, c). Although Group 1 and Group 2 regions showed opposite profiles regarding PU.1 and TCF1 co-binding, Group 3 regions displayed similar proportions of all possible combinations of PU.1 and/or TCF1 co-occupancies, and a minority with E proteins also (Fig. 7b, c).

Most partner factors were absent at the Group 4 OE-specific sites. PU.1 occupancy was detected in a minority of the open Group 4a sites, but was largely absent from the closed Group 4b sites. Furthermore, TCF1 did not actually occupy any Group 4 regions in normal Phase 2 pro-T cells (Fig. 7b), despite detectable TCF1 motifs in both Group 4a and Group 4b sites (Fig. 7a). As a result, the closed Group 4b sites could have isolated Runx binding without PU.1, TCF1, or E-protein engagement, unlike most natural Runx binding sites (Group 1, 2, 3). Unique among site Groups, Group 4b sites were also more highly associated with Runx non-DEGs than with Runx-responsive genes, indicating that they are probably functionally inert (Table S2)(Fig. S7a, b).

The sites most associated with Runx-activated or Runx-inhibited functional targets in these early Phase 1 cells were Group 3 sites, i.e., open and Runx-occupied in Phase 1 and Phase 2; Group 2a sites were also enriched near genes with Runx1 OE-dependent activation (Fig. S7a-c). Thus, Runx-induced factor TCF1 (encoded by *Tcf7*) could play a role in guiding Runx1 to Group 2 sites and/or increasing occupancy of Group 3 sites, to stimulate T-lineage progression. In accord, *Tcf7* knockdown using short hairpin RNA (shRNA) completely inhibited Bcl11b upregulation by Runx1 OE (Fig. 7e), suggesting that TCF1 is necessary for Runx factors to accelerate T-development.

### Combinatorial inputs from multiple TFs contributed to Runx-mediated gene regulation

To examine whether Runx factors also accelerate T cell development by a regulatory cascade through other TFs, we performed gene regulatory network inference analysis using Single Cell Regulatory Network Inference and Clustering (SCENIC)^60, 61^. SCENIC builds TF “regulons” by searching for co-expression of a TF and its potential regulatory target genes, then pruning the target list based on presence of putative regulators’ motifs within 10 kb of the TSS of each target gene. As the gene regulatory network is dynamically changing during T-developmental progression, we categorized cells based on their pseudotime value (Early ≤ 60, Mid 60-120, Late > 120) and compared enriched regulon activities between control, Runx1 OE, and Runx dKO cells (Fig. 8a) (Table S4). As expected, “Pseudotime Early” control cells showed strong regulon activities for Spi1, Irf8, and Mef2c, which were downregulated in “Pseudotime Late” control cells. However, these regulons remained strongly active in Runx dKO cells even after they reached “Pseudotime Mid”, demonstrating that the potential target genes of these factors depended upon Runx activity for their repression. Conversely, Runx1 OE cells had a shrunken Pseudotime Early category and showed much stronger regulon activities for Tcf7, Gata3, Patz1, Myb, Tcf12, Ets1, and Spib relative to control cells throughout Pseudotime Mid and Late, indicating that Runx-sensitive TFs enhanced both the onset and the magnitude of these regulon activities. Interestingly, some regulons such as Klf13 and Vezf1 were uniquely active in Runx1 OE cells, although these regulons were not dynamically activated in normal pro-T cells (Fig. 8b, c).

**Figure 8.**
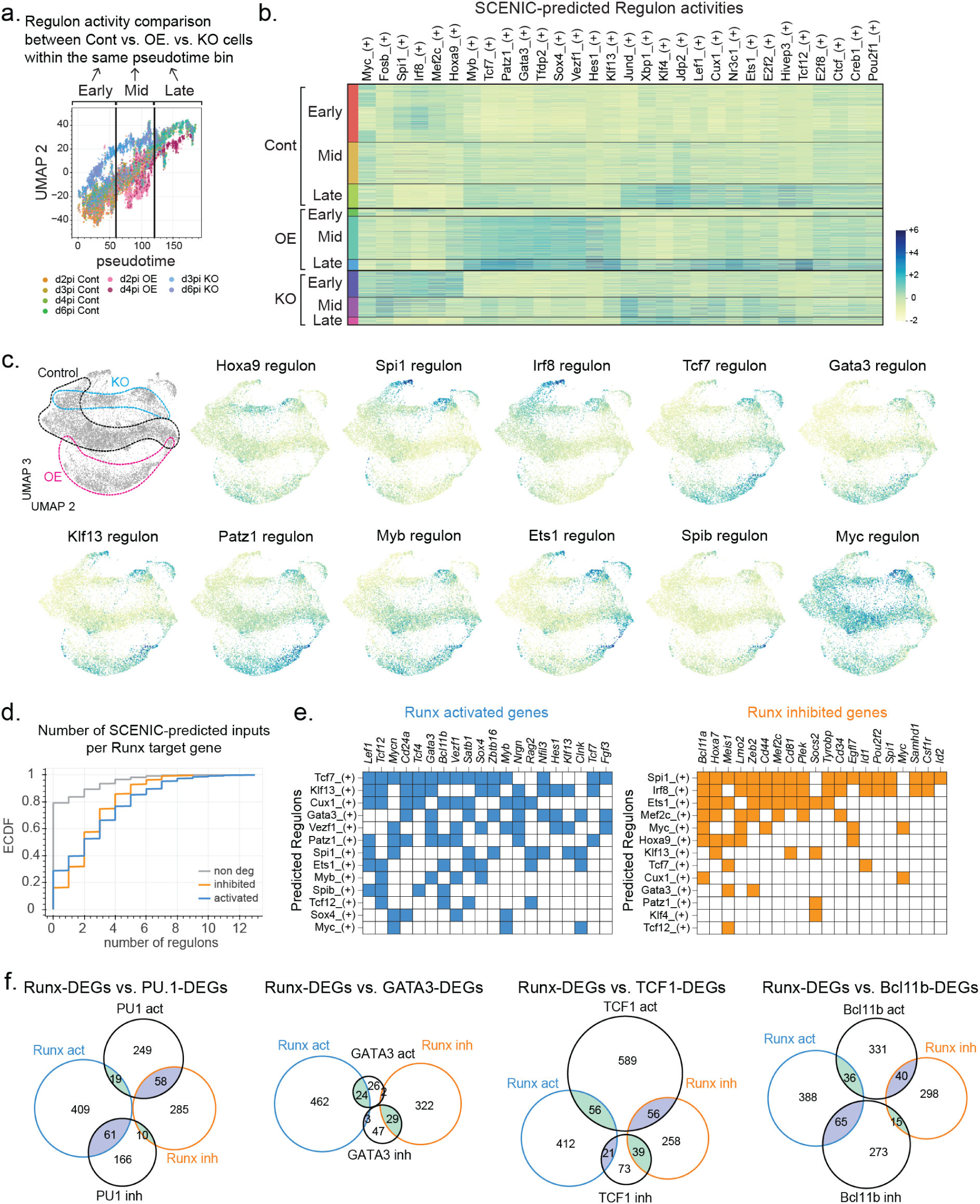
Runx TFs control gene regulatory network by cooperating with other TFs. **a,** Gene regulatory network analysis strategy is shown. Cells were grouped by Runx perturbation condition and pseudotime category to compute predicted target gene activity using SCENIC (pySCENIC, see Methods). **b-c,** SCENIC-predicted regulon activities for indicated TFs are represented as a heatmap (b) or highlighted on the UMAP2/UMAP3 manifold (c). The expressed regulons scoring adjusted p-value < 1e-10 from at least two different pairwise comparisons using Kolmogorov–Smirnov tests were selected to visualize. **d,** The members of each regulon were overlapped with Runx DEGs defined by KO and/or OE from Figure 3e. Then the numbers of overlapping predicted input regulons were enumerated per functionally responding Runx target gene or per non-DEG, and the results displayed as cumulative density functions. Kolmogorov–Smirnov test p-values were calculated by comparing Runx-activated or Runx-repressed DEGs with non-DEGs. Activated genes’ p-value=1.55e-15, inhibited genes’ p-value=8.88e-16. **e,** Curated Runx DEGs regulon memberships predicting input relationships are displayed as matrices. Colored cells in matrix indicate that a given Runx DEG (rows) is a member of a given regulon (columns). Blue; Runx-activated genes, orange; Runx-inhibited genes. **f,** Area-proportional Venn diagrams display overlap between functionally responsive Runx target genes with previously determined functional target genes of PU.1^13^, GATA3^55^, TCF1^55^, and Bcl11b^32^.

Next, we evaluated whether different regulon activities affected by Runx perturbation contributed to regulation of Runx target genes themselves. Importantly, 70-80% of Runx DEGs possessed at least one SCENIC-predicted input from a Runx-sensitive regulon, whereas 80% of Runx non-DEGs failed to overlap with any members of Runx-sensitive regulons (putative target genes). The enrichment of Runx-sensitive regulon membership among Runx DEGs suggests that Runx-dependent TF changes could significantly contribute to Runx impacts on target genes (Fig. 8d). Furthermore, 50% of Runx-activated and -inhibited target genes were predicted to be controlled by more than three SCENIC inputs, with each target gene predicted to receive different combinatorial inputs. For instance, *Lef1*, *Tcf12,* and *Mycn* (Runx-activated genes) were putative targets of 6-7 different regulators, with one common input (Tcf7) and 5-6 target-specific inputs. A similar trend was observed for Runx-inhibited genes, but receiving inputs from different sets of regulators.

The SCENIC analysis results were supported by Runx DEG overlap patterns with previously defined PU.1, TCF1, GATA3, and Bcl11b target genes (see Methods; Fig. S7d-g for gene names, Table S5 for full lists). Runx TFs mainly opposed PU.1 actions, as they had mostly opposite effects on the same genes (Fig 8f, S7d). In contrast, effects of GATA3 were strongly concordant with Runx responses of the same genes, including key molecules supporting T-developmental progression (Fig. 8f, S7e). Runx factors also worked with TCF1 to support T-cell identity and common lymphoid programs (Fig 8f, S7f). However, many TCF1-activated genes associated with cytokine response and proliferation were not co-activated by Runx factors, and both concordant and opposing responses were seen (Fig 8f, S7f). Finally, Bcl11b and Runx both supported T-identity associated program genes (Fig. 8f, Fig. S7g), but Bcl11b specifically repressed Runx1 OE-induced common lymphoid genes (Fig. S7g, left), thus presumably working to block access to ILC and NK cell potential. Together, these data showed a modular structure for the gene regulatory network that pro-T cells employ, in which Runx factors work as gene network mediators to oppose stem/progenitor/myeloid programming, while fueling common lymphoid and T-identity modules.

## Discussion

To achieve highly specific target gene regulation, regulatory elements in large metazoan genomes often exploit suboptimal binding affinities of TFs to enforce requirements for factor-factor collaboration^62–67^. Therefore, TF concentrations are an important parameter affecting TF occupancy at a particular DNA binding locus^68^. Runx factors can exert distinct developmental effects based on binding affinities^27^ and dose-dependent effects on hematopoietic progenitor emergence^69, 70^. We here show that dosage-sensitive DNA binding site choices by Runx TFs also had significant biological consequences during T-lineage specification. As chromatin states changed during T-lineage commitment, Runx1 binding remained within active chromatin compartments whether at normal or elevated factor levels. However, medium-quality Runx binding sites often recruited Runx factors stage-specifically, usually independent of chromatin state changes but utilizing functional collaborators. PU.1 appeared to be the main Phase 1 partner, with TCF1 and bHLH E proteins among others in Phase 2. This co-factor-associated Runx binding shift was sensitive to Runx availability, consistent with distinct partners competing for limited amounts of Runx factors. Thus, a modest increase in Runx levels in pre-commitment cells enabled Runx to bind precociously to post-commitment-specific Phase 2 sites, while still binding to Phase 1 sites.

Increased Runx availability concomitantly caused striking T-lineage developmental acceleration from DN1 at least to DN4 stage. This faster developmental progression was fueled by selective Runx activities upregulating common innate-lymphoid and T-identity programs driving Phase 1 to Phase 2 transition, before fully inducing cytokine and environment-responsive genes or completely shutting off Phase 1-associated genes. Notably, Runx1 OE-activated target genes were not only specific to T-cell identity but included genes that could drive ILC or NK development, though not TCR-γ locus transcripts (cf. ref. ^71^). The co-expression of Phase 1- and Phase 2-signature genes and incoherent activation of post-commitment programs suggest a modular structure for the pro-T cell gene regulatory network, in which Runx TFs minimally contributed to proliferation and cytokine-responses, while actively participating in T and innate-like cell-identity programs.

Reciprocal regulation by *Runx1/Runx3* dKO and by Runx1 OE highlighted the core Runx dose-dependent target genes. Added Runx occupancy upon OE was more associated with activated targets than with repressed targets. These Runx regulated genes included many TFs, which could contribute to the transcriptional profile both by co-binding with Runx1 to subsets of Phase 2-occupied sites, and by separate gene network effects. Most Runx DEGs were predicted to be co-regulated by these Runx-target TFs, and previously defined target genes of PU.1, GATA3, TCF1, and Bcl11b overlapped substantially with Runx-regulated genes. PU.1 and Runx activities largely opposed each other, whereas GATA3 and Runx effects were mostly concordant. Interestingly, TCF1 and Runx TFs collaboratively supported T cell/ ILC identity programs, but genes related to cellular proliferation, metabolism, and cytokine responses were not co-regulated by these factors. Finally, while both Bcl11b and Runx provided positive inputs to T-identity related genes, Bcl11b inhibited the innate-like program genes that were induced by Runx1 OE. Thus, Runx factors function as dose-dependent gene network mediators to orchestrate discrete transcriptome modules during early T cell development.

## Acknowledgments

We thank Rothenberg lab members for helpful discussions, Rochelle Diamond and members of the Caltech Flow Cytometry and Cell Sorting facility for sorting, Igor Antoshechkin and Vijaya Kumar of the Caltech Jacobs Genomics Facility for bulk RNA sequencing, Henry Amrhein and Diane Trout for computer support, Jeff Park and Sisi Chen from the Caltech Single Cell Profiling and Engineering Center for providing support for processing 10X Chromium samples, Ingrid Soto for mouse care, Maria Quiloan for mouse genotyping and supervision.

## Data and Materials availability

All genomic sequencing data have been deposited in Gene Expression Omnibus under accession numbers GSE218147 (C&R and ChIP-seq) and GSE218149 (scRNA-seq). All other data needed to evaluate the conclusions in the paper are present in the paper or the Supplementary Materials.

## Author contribution

B.S. and E.V.R. conceptualized the project, wrote the paper, and edited the paper. B.S. performed the experiments, and analyzed data with W.Z. and J.W. F.G. wrote the in-house bioinformatic pipeline for hashtag alignment and provided further analysis. E.V.R. supervised research, acquired funding, and provided additional data analysis. All authors edited the paper and provided helpful comments.

## SUPPLEMENTAL MATERIALS

### Supplementary Tables

**Table S1. Runx sensitive genes defined by scRNA-seq.**

Differentially expressed genes in control vs. *Runx1/Runx3* dKO or control vs. Runx1 OE groups are shown. “Runx Core DEGs” marks whether a given gene is sensitive to both Runx dKO and Runx1 OE. “Category” indicates whether a give gene responded to Runx dKO and/or OE. “dn” means downregulated in comparison to control cells and “up” means upregulated in comparison to control cells. “Developmentally dynamic category” is determined by differential gene expression analysis comparing cells in cluster 2 (early) vs. cluster 1 (late). If a gene is significantly highly expressed in cluster 2, the gene is marked as Phase 1 DEG. If a gene is significantly upregulated in cluster 1, the gene is annotated as Phase 2 DEG.

**Table S2. scRNA-seq vs. bulk-RNA-seq comparison and Runx occupancy annotation.**

Differentially expressed gene (DEG)s determined by scRNA-seq and previously reported bulk-RNA-seq data were annotated. Previously reported bulk-RNA-seq included two different timepoints of *Runx1/Runx3* dKO. “Phase 1 bulk” was measured by introducing g*Runx1/Runx3* before T-lineage commitment for 3 days (identical to the d3 post-infection timepoint in this study). “Phase2 bulk” was measured by deleting *Runx1/Runx3* after 10 days of OP9-Dll1 co-culture (post-commitment) by introducing gRNA for 3 days. If a gene is scored as a DEG by an indicated method and timepoint, it was marked as “1”; if a gene was categorized as a non-DEG, it was marked as “0”. In addition, the numbers of annotated non-promoter Runx peaks in the indicated group (Group 1, Group 2a, Group 2b, Group 3, Group 4a, and Group 4b) for each gene are marked.

**Table S3. Total list of differentially expressed genes in “OE-accelerated UMAP-2 window” from Figure 4.**

Differentially expressed genes in control cells vs. Runx1 OE cells within in the UMAP 2 values ranging from −30 to 5 are listed. Average Log_2_FC is calculated by comparing control / OE cells (genes expressed highly in control cells are positive).

**Table S4. SCENIC predicted Runx regulon members and their overlaps with Runx target genes.**

The putative target genes (regulon members) predicted by SCENIC in the scRNA-seq dataset are listed. The KS test p-values for indicated comparisons are shown. The overlap between Runx target genes and predicted regulon members are marked.

**Table S5. Comparison between Runx sensitive genes and other TF-regulated genes presented in Figure 8.**

Runx DEGs defined by Runx1 OE and/or *Runx1/Runx3* dKO were compared with the genes activated or inhibited by the indicated TF that were previously reported.

**Supplementary figure 1. Distinct motif enrichment patterns in dynamically shifting Runx binding sites, comparison with PU.1 site stability, and efficient detection of direct Runx binding sites by CUT&RUN.**

**a,** Heatmap illustrates PU.1 binding profiles in immortalized HSPC, DN1, DN2a, and DN2b cells. **b,** Top motifs enriched in Runx binding sites from indicated regions of Figure 1b are shown. **c,** Scatter plots and Area-proportional Venn diagrams compare Runx1 and Runx3 binding sites in Phase 1 and Phase 2 pro-T cells, as measured by ChIP-seq cross-linked with DSG+FA vs. by CUT&RUN (C&R). Numbers in the Venn diagram indicate number of differential peaks compared between DSG-crosslinked ChIP-seq vs. C&R (fold enrichment>2, Poisson enrichment p-value<0.001). **d,** Violin plots show Runx motif quality position weight matrix (PWM) score in non-promoter Runx peaks detected similarly by ChIP-seq and C&R (purple) or preferentially detected by different technique (green; more efficiently detected by C&R, red; more efficiently detected by ChIP-seq). The horizontal dotted black line shows threshold PWM score to be recognized as a Runx motif. Thin vertical black lines mark 1.5x interquartile range and thick vertical black lines indicate interquartile range. The red lines with white circles show median values. **e,** Motif frequencies for Runx, bHLH, ETS, and PU.1 or TCF1 (Tcf7) factors within each peak are displayed. **f,** Percentages of ChIP-seq or C&R-detected Runx binding sites that are open or closed at a given stage is shown. Only non-promoter sites were calculated as most of the promoter sites are stably accessible.

**Supplementary figure 2. Runx TFs predominantly interact with active large-scale chromatin compartments, yet local chromatin state is not a major barrier for stage-specific redeployment of Runx factors.**

**a-b,** Genomic regions were assigned to compartment A (active, HiC PC1 value ≥ 10), compartment B (inactive, HiC PC1 value ≤ -10), and compartment N (neutral, -10 < HiC PC1 value < 10) in 1kb-bins from DN1 (ETP), DN2, and DN3 cells (data from ref. ^22^). Then the regions stably maintaining compartment states (A-to-A or B-to-B) vs. the regions undergoing compartment flipping were categorized and their enrichment within different groups of Runx binding sites or total genomic regions were compared. Graphs in inset show expanded-scale view from Fig S2a to record rare changes in genomic compartment reprogramming during DN1 (ETP) to DN3 progression. **b,** Representative UCSC genome browser tracks near *Bcl11b* and *Ets1* regions show compartment state (represented with HiC PC1 values), DN1 and DN3 Runx occupancies, and published ATAC signals with CTCF and SMC3 ChIP-seq. **c,** Heatmaps represent distinct chromatin states computed using ChromHMM. The enrichments of different histone marks, ATAC, and loop-forming machineries (CTCF, SMC3) with each chromatin state are shown in purple (left). Genomic annotation for chromatin states is displayed in orange (middle). Enrichment with different groups of Runx peaks is illustrated in blue (right). Constitutively occupied Group 3 peaks and promoter peaks were enriched among constitutively active chromatin states (states 8 and 9), as expected, and Group 1 peaks (losing Runx binding from Phase 1 to Phase 2) had the highest enrichment within Phase 1-preferential active states (states 1 and 2). In contrast, however, the Group 2 sites newly occupied during commitment were more enriched among constantly accessible regions with weak H3K4me2 marks (state 10), even more than they were enriched for Phase 2-specific active states (states 5 and 6). Constitutively active states (states 8 and 9) were also enriched for Group 2 peaks, and Group 2 peaks were also the only group showing enrichment among sites that were largely ATAC-closed in both Phase 1 and Phase 2 (state 7).

**Supplementary figure 3. Increase in Runx1 availability prevents PU.1-mediated Runx1 depletion.**

**a,** Experimental design to test Runx-dosage and co-factor dependent Runx redistribution in DN3-like Scid.adh.2C2 cells. **b,** Histograms show protein expression levels of PU.1 and Runx1 after introducing PU.1 and/or Runx1-expressing vectors. Bar graphs summarize geometric mean fluorescent intensities (gMFI) of PU.1 and Runx1 with means and standard deviations. 6 independent experiments. **c,** Expression of non-T-lineage markers, CD11b and CD11c, was measured using flow cytometry. Bar graphs show frequencies of cells that do not express these markers. Mean and standard deviation from 6 independent experiments are displayed. One-way ANOVA. **d,** Peak-centered heatmap illustrates Runx1 and PU.1 binding patterns in non-promoter sites under indicated conditions from 2 independent ChIP-seq experiments. **e,** Density plots display motif frequencies for Runx1 and PU.1 in each peak and violin plots illustrate the best motif qualities for Runx1 and PU.1 in a given peak. **f,** Representative histograms display Runx1 expression levels at 4 days after transducing Runx1-OE or empty-control vectors. Cells were gated on live alternative lineage^-^ infection^+^ cells, then separated as cKit^hi^ CD25^-^ (DN1) and cKit^hi^ CD25^+^ (DN2a) populations. Graph summarizes results from 7 independent experiments. Two-way ANOVA. ***=p-value<0.001, **=p-value<0.01.

**Supplementary figure 4. Single-cell transcriptome analyses of Runx perturbations: deviations from normal developmental clusters due to effects on core target genes responding to both gain- and loss-of-functions.**

**a-b,** tSNE1-2 (a) and UMAP1-2 (b) display transcriptomes of control- or Runx1 overexpressed (OE) or *Runx1/Runx3* knockout (KO) cells at indicated timepoints (left). Genes associated with different stages of cell cycles are illustrated on tSNE 1-2 (right). Top panels show location of cells before cell-cycle regression and bottom panels illustrates distribution of cells after cell-cycle regression. Note that Runx1 OE tends to shift population toward G1 while KO shifts cells toward G2/M, but Runx perturbation states do not separate well on these axes. **c,** Cluster distributions of indicated Runx- perturbation conditions are shown. Size of each dot represents number of cells and colormap indicates z-score from standard residual analysis followed by Fisher’s exact test. **d,** Expression patterns of stem or myeloid-associated genes, *Cd81*, *Csf2b*, *Meis1*, and *Ifngr2* are displayed on UMAP2-3 axes. **e,** Scatter plots compare Log_2_ fold-changes (FC) in gene expression between Runx1 OE vs. control or Runx KO vs. control populations at different timepoints (d2 vs. d4 after introducing OE conditions, d3 vs. d5 after delivering gRNA for KO conditions). Each dot represents a different gene. **f,** Heatmap illustrates expression profiles of the common Runx target genes sensitively responding to both Runx1 OE and Runx KO. Each cluster is sorted by developmental progression order.

**Supplementary figure 5. Runx1 overexpression inhibited myeloid and granulocyte program, while supporting NK cell program even after inducing Bcl11b expression**

**a,** Schematics illustrate experimental design for competitive commitment assay. Empty control or Runx1 overexpression vectors expressing different markers were cultured with OP9-Dll1 to initiate T-cell development. After 2 days, DN1, Bcl11b^-^DN2a, and Bcl11b^+^ DN2a cells were each sorted from each condition. The same number (100 cells) of the same stage cells from control and Runx1 OE conditions were co-cultured with Notch-signaling (OP9-Dll1) or Notch nonsignaling (OP9-Control) stromal cells for 6 days, supplemented with IL-7 and Flt3-ligand. **b,** Representative flow plots show competition outcomes between Control (x-axis) vs. Runx1 OE (y-axis) from each condition. Graphs summarize the absolute number and frequencies of control vs. Runx1-OE populations in both conditions. Runx1 OE cells were disfavored with and without Notch signaling. **c,** Expression of NK1.1 vs. Ly6G/Ly6C were measured by flow cytometry after culture without Notch signals. Graphs show frequencies of cells expressing Ly6G/Ly6C or NK1.1 in cells derived from the indicated input populations. 2 independent experiments, n=8-10, Two-way ANOVA. ***=p-value<0.001, **=p-value<0.01, ns=not significant.

**Supplementary figure 6. Elevated Runx1 levels in Phase 1 resulted in additional Runx occupancies in post-commitment preferred sites and closed chromatin regions**

**a,** Gating strategy to sort Phase 1 cells for C&R is illustrated. Briefly, bone-marrow progenitor cells were co-cultured with OP9-Dll1 cells for 2 days, and empty control or Runx1 overexpressing vector was retrovirally introduced. After 40-42 hours (total 4 days of culture on OP9-Dll1 cells), infection^+^ Phase 1 cells were sorted. In this system, for most cells to reach Phase 2 normally, 8-10 days of culture are needed^1, 72^. **b,** Scatter plots and Venn diagrams compare differential Runx1 occupancies at promoter (top) and non-promoter regions (bottom) when Runx1 concentration was increased. Numbers indicate differential Runx1 binding sites (fold enrichment > 2, Poisson enrichment p-value<0.001). **c,** Runx1 C&R signal intensities from indicated cells are shown. Note increased occupancy even at Group 1 and Group 3 sites which were already bound in control Phase 1 cells. **d,** UCSC genome browser tracks show Runx binding patterns (orange tracks, experimental conditions; blue tracks, unperturbed pro-T cells), PU.1 in DN1 cells, TCF1 in DN3 cells, E2A and HEB in DN3 cells, and ATAC-seq signals (black) in Phase 1 (DN1) and Phase 2 (DN2b) cells. Regulatory regions for *Plek, Lmo2, Meis1, Cd3* clusters, *Tcf7*, and *Thy1* are shown. **e,** Bar graph represents compartment state profiles within different groups of Runx binding sites.

**Supplementary figure 7. Runx TFs show distinct regulatory relationships with PU.1, GATA3, TCF1, and Bcl11b for different gene regulatory modules.**

**a,** Specific associations between different classes of Runx binding sites and Runx DEGs are tested using Fisher’s exact test. Graphs visualize the percentages of the genes associated with such peaks (height of the spike) and the number of genes possessing at least one Runx binding in DEG groups (size of hexagon). Gray bars to the left of each plot indicate the percentages of genes associated with each peak type among non-responding DEGs, and broken line uses this level as a reference for DEG enrichment. All site types except Group 4b sites were significantly enriched among DEGs relative to non-DEGs (shown by relatively higher of spike heights compared to non-DEGs). Color map compares particular types of response to Runx perturbation as compared to other responses to perturbation, among the DEGs with a given site type. Colors depict z-scores (standardized residuals), calculated for relative enrichment of a given association within the DEG groups. For example, dark cyan indicates that genes linked to a given site Group are especially positively enriched for the indicated response type. See Methods for how the non-DEGs and the core DEGs were defined. *p-value<0.05, **p-value<0.01, ***p-value<0.001. **b,** Association of different groups of Runx binding sites with Runx target genes are shown. Violin plots show percent of each group of Runx peaks among total number of Runx peaks in a given gene. **c,** Diagrams show a schematic summary of different groups of Runx peaks found commonly near Runx DEGs and Runx non-DEGs (Runx-independent). Note that each gene can possess multiple types of Runx peaks. **d-g,** Area-proportional Venn diagrams display overlap patterns found between Runx DEGs with previously characterized functional targets of the indicated TFs. For Runx DEGs, genes activated (blue) or inhibited (orange) by Runx1 OE vs. Runx KO are each shown. Informative genes overlapping different classes of functionally responsive Runx DEGs are listed in different colored fonts: overlaps with Core-responsive DEGs showing reciprocal effects of Runx1 OE and KO (red); overlaps with DEGs defined by Runx1 OE-responses only (green); and overlaps with DEGs defined by Runx KO-responses only (blue) are listed. Comparisons between **d,** PU.1 target genes, **e,** GATA3 target genes, **f,** TCF1 target genes, and **g,** Bcl11b target genes are shown.

## NATURE METHODS

### Animal studies

C57BL/6J (B6), B6.Cg-Tg(BCL2)25Wehi/J (Bcl2-tg), B6.Gt(ROSA)26Sortm1.1(CAG-cas9*,-EGFP)Fezh/J (Cas9) mice were purchased from the Jackson Laboratory (#000664, #002320, #026179) and bred at the California Institute of Technology. B6.Bcl11b^mCitrine/mCitrine^ (B6.Bcl11b-mCitrine reporter) mice were described previously ^49, 73^. Both male and female mice were used for this study. All animals were bred and maintained under specific pathogen-free conditions at the California Institute of Technology according to Institutional Animal Care and Use Committee (IACUC) regulations.

### Cell Lines

The OP9-Dll1 (obtained from Dr. J. C. Zúñiga-Pfl^ü^ cker^48^) or mouse MS4-Dll4 (obtained from Dr. Gay Crooks^59^) stromal cell lines were utilized for *in vitro* cell culture to recapitulate early thymic T-cell development. The stromal cell lines were maintained as described in the original references. The Scid.adh.2c2 DN3-like cell line^74^ was cultured in RPMI1640 with 10% fetal bovine serum, 2 mM glutamine, 100 IU/mL penicillin, 100 μg/mL streptomycin, 0.1 mM sodium pyruvate, non-essential amino acids, and 50 μM β-mercaptoethanol.

### *In vitro* OP9 cell culture

Bone marrow was obtained from the femurs and tibiae of 8-12 week-old B6. Bcl2-tg or progeny of B6.Bcl2-tg x Bcl11b^mCitrine/mCitrine^ or progeny of B6.Cas9 x Bcl2-tg mice. The The *Bcl2* transgene supports improved cell recovery under regulatory perturbation without altering development^75–77^. Progenitor cells from the bone marrow cell suspension were enriched by depleting mature lineage^+^ cells expressing CD3ɛ (clone 145-2C11), CD19 (clone 1D3), B220 (clone RA3-6B2), NK1.1 (clone PK136), CD11b (clone M1/70). CD11c (clone N418), Ly6G/C (clone RB6-8C5), and Ter119 (clone TER-119) using MACS LS magnetic columns (Miltenyi Biotec). Enriched progenitor cells were co-cultured with OP9-Dll1 cells and supplemented with 1 ng/mL of IL-7 (Peprotech) and 10 ng/mL of Flt3L (Peprotech) in OP9 medium (α-MEM, 20% FBS, 2 mM glutamine, 100 IU/mL penicillin, 100 mg/mL streptomycin, and 50 µM β-ME). OP9 *in vitro* cultures were done under 37 °C, 7% CO_2_ environment.

To obtain unperturbed Phase 1 pro-T cells for CUT&RUN, bone-marrow progenitor cells were cultured with OP9-Dll1 cells for 5 days with 10 ng/mL IL-7 and Flt3L each. To measure Runx1 binding sites after retroviral infection, bone-marrow progenitor cells were cultured on OP9-Dll1 cells for 2 days and either empty control vector or Runx1 overexpression vector was introduced for 40-42 hours.

### Mixed chimeric Artificial Thymic Organoid (ATO) 3D culture

Bone-marrow progenitor cells obtained as described above were incubated with 10 ng/mL IL-7, 10 ng/mL of Flt3L, and 10 ng/mL of SCF in OP9 medium overnight to launch the cells into cycle. Then progenitor cells were infected with control or Runx1 overexpressing MSCV vector expressing mCherry or human NGFR marker and incubated with 10 ng/mL IL-7, 10 ng/mL of Flt3l, and 0.1 ng/mL SCF in OP9 medium (SCF concentration was reduced to recover surface cKit expression). After 24 hours delivering retroviral vector, infection marker^+^ lineage (TCRβ, TCRγδ, CD19, NK1.1, CD49b, Ly6G/C, CD11b, CD11c)^-^ Sca1^+^ cKit^+^ (LSK) cells were FACS sorted. The ATOs were formed and maintained by following the original reference^59^. Briefly, 1,000 of each infection marker^+^ LSK cells and 150,000 mouse MS4-Dll4 cells were aggregated and seated at the air-medium interface on a culture insert (Millipore Sigma) in serum-free ATO medium (DMEM-F12, 1X B27, 2 mM glutamine, 100 IU/mL penicillin, 100 μg/mL streptomycin, 30 μM Ascorbic acid) supplemented with 5 ng/mL of IL-7 and 5 ng/mL of Flt3L. The cytokine-supplemented culture medium was replaced every 3 days and IL-7 and Flt3L concentrations were dropped to 1 ng/mL (each) after day 10.

### Retroviral transduction

Mouse Runx1 full-length sequence was inserted into the murine stem cell virus (MSCV) retroviral-mCherry or MSCV-human NGFR vector (Addgene #80157, 80139) as previously described^49, 71^. The guide-RNA (gRNA) against Runx1 or Runx3 were inserted into E42-human NGFR or E42-mTurquoise2 vector as previously described^1, 13^. Three gRNAs were utilized to target each Runx paralog (Addgene #189799, #189800, #189801, #189802, #189803, #189804, #189805, #189806). For retroviral infection, the target cells were centrifuged at 500×g, 32°C for 2 hours with viral supernatant supplemented with 8 µg/mL polybrene. After the spinfection, viral supernatant was removed and replaced with cytokine-supplemented culture medium.

### Flow cytometry analysis and cell sorting

Cell surface staining was performed following Fc blocking by incubating single cell suspensions in 2.4G2 hybridoma cell supernatant. Then cells were stained with a biotin-conjugated lineage cocktail: TCRβ (BioLegend, clone H57-597), TCRγδ (eBioscience, clone GL-3), CD19 (BioLegend, clone 6D5), NK1.1 (BioLegend, clone PK136), CD49b (BioLegend, HMa2), CD11b (BioLegend, clone M1/70), CD11c (BioLegend, clone N418), and Ly6G/C (BioLegend, clone RB6-8C5). Secondary surface staining was performed with fluorescently conjugated streptavidin, CD45 (eBioscience, clone 30-F11), cKit (eBioscience, clone 2B8), CD44 (eBioscience, clone IM7), CD25 (eBioscience, clone PC61.5), and hNGFR (BioLegend, clone ME20.4). A viability dye (Life Technologies, Aqua) or 7AAD (eBioscience) was applied to exclude dead cells.

For intracellular staining of TFs, cells were fixed with 2% paraformaldehyde for 15 min at room temperature after surface staining. Then cells were permeabilized with the Foxp3 Permeabilization/Fixation kit (eBioscience) and stained with fluorescently conjugated antibody against Runx1(eBioscience, clone RXDMC), against TCF1 (CST, clone C63D9), against GATA3 (BD, clone L50-823), or against PU.1 (CST, clone 9G7), or an isotype control (BioLegend, clone RTK2758). Samples were acquired using a CytoFlex analyzer (Beckman Coulter) and data was analyzed with FlowJo v.10.8.1 (BD). Except for lineage commitment assay (Fig. S5), cells were gated on live, alternative lineage (TCRβ, TCRγδ, CD19, NK1.1, CD49b, CD11b, CD11c, Ly6G/C)^-^ infection^+^ CD45^+^ population for analysis.

For single cell RNA-seq, bone marrow progenitor cells were subjected to *in vitro* culture as described. On day 2, 3, 4, or 6 post infection, Phase 1 cells from each experimental condition were stained with unique hashtag-oligo (HTO) antibody (BioLegend, TotalseqA

HTO1-HTO8), then cells were sorted for Lineage^-^ CD45^+^ cKit^high^ mCherry^+^ (marker for MSCV vector) or mTurquoise2^+^ hNGFR^+^ (markers for gRNA expressing vectors) population using BD FACSAria Fusion at the California Institute of Technology Flow Cytometry Facility.

The following conditions were subjected to each scRNA-seq experiment.

Experiment 1:

1. day 2 post-infection MSCV control rep 1
2. day 2 post-infection MSCV control rep 2
3. day 2 post-infection MSCV Runx1 OE rep 1
4. day 2 post-infection MSCV Runx1 OE rep 2
5. day 3 post-infection gRNA control rep 1
6. day 3 post-infection gRNA control rep 2
7. day 3 post-infection gRNA Runx1/Runx3 dKO rep 1
8. day 3 post-infection gRNA Runx1/Runx3 dKO rep 2

Experiment 2:

1. day 2 post-infection MSCV control rep 3
2. day 4 post-infection MSCV control
3. day 2 post-infection MSCV Runx1 OE rep 3
4. day 4 post-infection MSCV Runx1 OE
5. day 3 post-infection gRNA control rep 3
6. day 6 post-infection gRNA control
7. day 3 post-infection gRNA Runx1/Runx3 dKO rep 3
8. day 6 post-infection gRNA Runx1/Runx3 dKO

### CUT&RUN (C&R)

C&R was performed by following original methods previously described^39, 40, 78^ with minor modifications. Briefly, pro-T cells were FACS sorted and washed with wash buffer (100 mM NaCl, 20 mM HEPES, 0.5 mM spermidine, 1X protease inhibitor, and 0.5% BSA) twice. Then, 400-500K DN3 cells were bound to 20 μL of activated concanavalin-A coated beads by incubating in wash buffer at room temperature for 5-10 min. For Phase 1 pro-T cells obtained from OP9-Dll1 culture (4-5 days of culture), 180-250K cells were used and cells were bound to 10 μL of activated concanavalin-A coated beads. The bead-bound cells were incubated with anti-rabbit antibodies for Runx1 (abcam, ab23980), Runx3 (gift from Dr. Yoram Groner^79^), TCF1(CST 2203, CST 2206, ab30961, ab183862, note that ab183862 was discontinued due to cross-reactivity between different members of the TCF7 family), E2A (abcam, ab228699), HEB (Proteintech 14419-1-AP) or negative control antibody (guinea pig anti-rabbit antibody, Antibodies-Online, ABIN101961). Cells were incubated in 100 μL (180-250K cells) or 200 μL (400-500K cells) of antibody buffer (0.0005-0.001% wt/vol digitonin in wash buffer with 1 mM EGTA, 1-2 μg antibody) for 2 hours at 4 °C. After antibody incubation, permeabilized cells were washed with digitonin buffer (0.0005-0.001% wt/vol digitonin in wash buffer) and incubated with 700 ng/mL of protein A-MNase (pA-MN) in a total volume of 250 μL for 1 hour at 4 °C. For chromatin digestion of thymic DN3 cells, cells were incubated with 2 mM CaCl_2_ in 150 μl digitonin buffer at 0 °C for 30 min, and the reaction was stopped by adding 100 μL 2X stop buffer (340 mM NaCl, 20 mM EDTA, 4 mM EGTA, 100 μg/mL RNase A, 50 μg/mL glycogen, 0.0005-0.001% digitonin). For chromatin digestion of Phase 1 cells, cells were washed with low-salt rinse buffer (0.5 mM spermidine, 20 mM HEPES, 0.0005-0.001% digitonin, 1x protease inhibitor) and incubated with 200 μL of low-salt high-Ca^2+^ incubation buffer (3.5 mM HEPES, 10 mM CaCl_2_, 0.0005-0.001% digitonin) at 0 °C for 5 min. After digestion, incubation buffer was quickly replaced with 200 μL of 1X STOP buffer (170 mM NaCl, 20 mM EGTA, 50 μg/mL RNAse, 25 μg/mL glycogen). Digested chromatin was released by incubating at 37 °C for 15 min and centrifuged at 4 °C at 16000g for 5 min. DNA was extracted by incubating with 0.1% SDS and 20 mg/mL of Proteinase K at 50 °C for 1 hour, followed by Phenol Chloroform extraction.

### PU.1 titration by Runx1 addition

ChIP-seq was performed as described previously^13, 32, 53^. Briefly, Scid.adh.2c2 cells were infected with pMXs-PU.1-human NGFR or pMXs-control-human NGFR vector in combination with MSCV-Runx1-HA-mCherry or MSCV-control-mCherry vector. At day2 post-infection, NGFR^+^ cells were enriched using MACS LS magnetic columns and 7×10^6^of NGFR^+^ Scid.adh.2c2 cells were crosslinked with 1mg/mL DSG (Thermo Scientific) followed by 1% formaldehyde. The reaction was quenched by 0.125M glycine. Nuclei were isolated by incubating crosslinked cells in Nuclei Isolation buffer (50 mM Tris-pH 8.0, 60 mM KCl, 0.5% NP40) and lysed in Lysis buffer (0.5% SDS, 10 mM EDTA, 0.5 mM EGTA, 50 mM Tris-HCl (pH 8)). The lysates were sonicated on a Bioruptor (Diagenode) for 18 cycles (one cycle: 30sec max power sonication followed by 30 sec rest). Rabbit anti-HA antibody (Santi Cruz) was bound to Dynabeads anti-Rabbit (Invitrogen) and incubated with sonicated chromatin in 1X RIPA buffer at 4°C overnight. After washes, precipitated chromatin fragments were eluted in ChIP elution buffer (20 mM Tris-HCl, pH 7.5, 5 mM EDTA 50 mM NaCl, 1% SDS, and 50 μg proteinase K) by incubating at 65°C for 14 hours. Eluted DNA was cleaned up using Zymo ChIP DNA Clean & Concentrator according to manufacturers’ protocols.

### Single cell RNA-seq

For single cell RNA-seq, pro-T cells obtained from OP9-Dll1 culture were stained with surface antibodies followed by hashtag oligo labeling with TotalSeq A (BioLegend) anti-Mouse Hashtag 1-8 (1:50, in separate samples). After FACS sorting the target cells, samples were washed with 1X HBSS supplemented with 10% FBS and 10 mM HEPES and resuspended to 1×10^6^ cells/1mL concentration. Then, 16,000 cells were loaded into a 10X Chromium v3 lane, and the subsequent preparation was conducted following the instruction manual of 10X Chromium v3.

### Library preparation and deep sequencing

C&R libraries were prepared using NEBNext ChIP-Seq Library Preparation Kit (NEB) by following previously published protocol^80^. ChIP-seq libraries were prepared using NEBNext ChIP-Seq Library Preparation Kit (NEB) according to the manufacturer’s protocol. Single cell RNA-seq cDNA libraries were prepared using 10X Chromium 3’ capture v3 kit. Single cell hashtag oligo library was prepared by following the BioLegend TotalseqA guide. After the library preparation, the sequencing was performed with paired-end sequencing of 50 bp (C&R and ChIP-seq) or 150 bp (single-cell RNA-seq) using HiSeq4000 by Fulgent Genetics, Inc. (Temple City, CA) or NextSeq by the California Institute of Technology Genomics core. Each ChIP-seq library was sequenced to a targeted depth of 30 million reads. For C&R libraries, each sample was sequenced to a targeted depth of 10 million reads. Single cell RNA-seq cDNA libraries were sequenced to a targeted depth of 65,000-70,000 reads per cell and hashtag oligo libraries were sequenced for 2,000-2,500 reads per cell.

### C&R, ChIP-seq, and ATAC-seq analyses

Sequenced reads from ChIP-seq and C&R libraries were mapped to the mouse reference genome GRCm38/mm10 using Bowtie2 (v3.5.1)^81^ and reproducible peak calling was performed using a HOMER (v.4.11.1)^82^ adaptation of the Irreproducibility Discovery Rate (IDR) tool according to ENCODE guideline. For downstream analysis, peaks with a normalized peak score ≥ 15 (for ChIP-seq) or peak score ≥ 10 (for C&R) were considered. Publicly available ATAC-seq data (GSE100738) was downloaded as raw sequence read files and mapped onto GRCm38/mm10. After filtering out mitochondrial reads using Samtools (v.1.9), peak calling was conducted with Genrich (v.0.6). Peaks were annotated to genomic regions using HOMER package (annotatePeaks.pl). Genes associated with C&R peaks were annotated using GREAT (v.4.0.4) with proximal: 5kb upstream, 1kb downstream, plus distal: up to 1000kb mode^83^. For UCSC Genome Browser visualization, bigwig files were generated from the aligned bam file using deepTools (bamCoverage -- binSize 20 --normalizeUsing CPM).

Differentially occupied peak analysis was performed using a HOMER package (getDifferentialPeaks.pl) and the resulted groups were visualized as heatmaps or area proportional Venn diagrams or scatter plots. Area proportional Venn diagrams were generated using Python Matplotlib-venn tools (v.0.11.7) or R eulerr package (v. 6.1.1). Scatter plots were generated by counting tag densities from indicated tag directories. The resulting tag counts per 10 million reads (base 2 logarithmic converted) were visualized using Python holoviews (v.1.15.0) with datashader (v.0.14.2) operation.

Peak centered heat maps were created with a deepTools2^84^ (v 3.5.1) in a 3000 bp region by computing matrix (computeMatrix reference-point --referencePoint center -b 1500 -a 1500 -R -S --skipZeros) and then visualized (plotHeatmap). In order to reference points for heat maps, co-occurring or unique peaks were computed using the HOMER package (mergePeaks -venn) and each cluster groups were defined by Boolean logic. Only non-promoter peaks were considered unless marked as “promoter”.

### Motif density, enrichment, and quality analyses

For quantitative analysis of motif frequencies, we used the HOMER package (findMotifGenome.pl) using a 200bp window and *De novo* results were utilized. We first examined whether the DNA sequences in each group were more or less favorable for recruiting Runx factors themselves, by analyzing the frequency of a Runx motif occurrence per peak (normalized to the peak size). The frequency of motif occurrence in regions of 2000 bp surrounding the C&R or ChIP-seq peak sites were analyzed by using a HOMER package (annotatePeaks.pl -size <#> -hist <#> -m < MOTIF file >). The resulting histograms were visualized using Python bokeh plotting (v.2.4.3).

The motif score results throughout this paper represent the best motif quality in the peak, based on position weight matrix (PWM, referred to as a motif score). The motif score of each peak was calculated as described previously^13^ using a HOMER function (annotatePeaks.pl -m <MOTIF file> -mscore).

### Published data used

Following publicly available data are utilized for analysis:

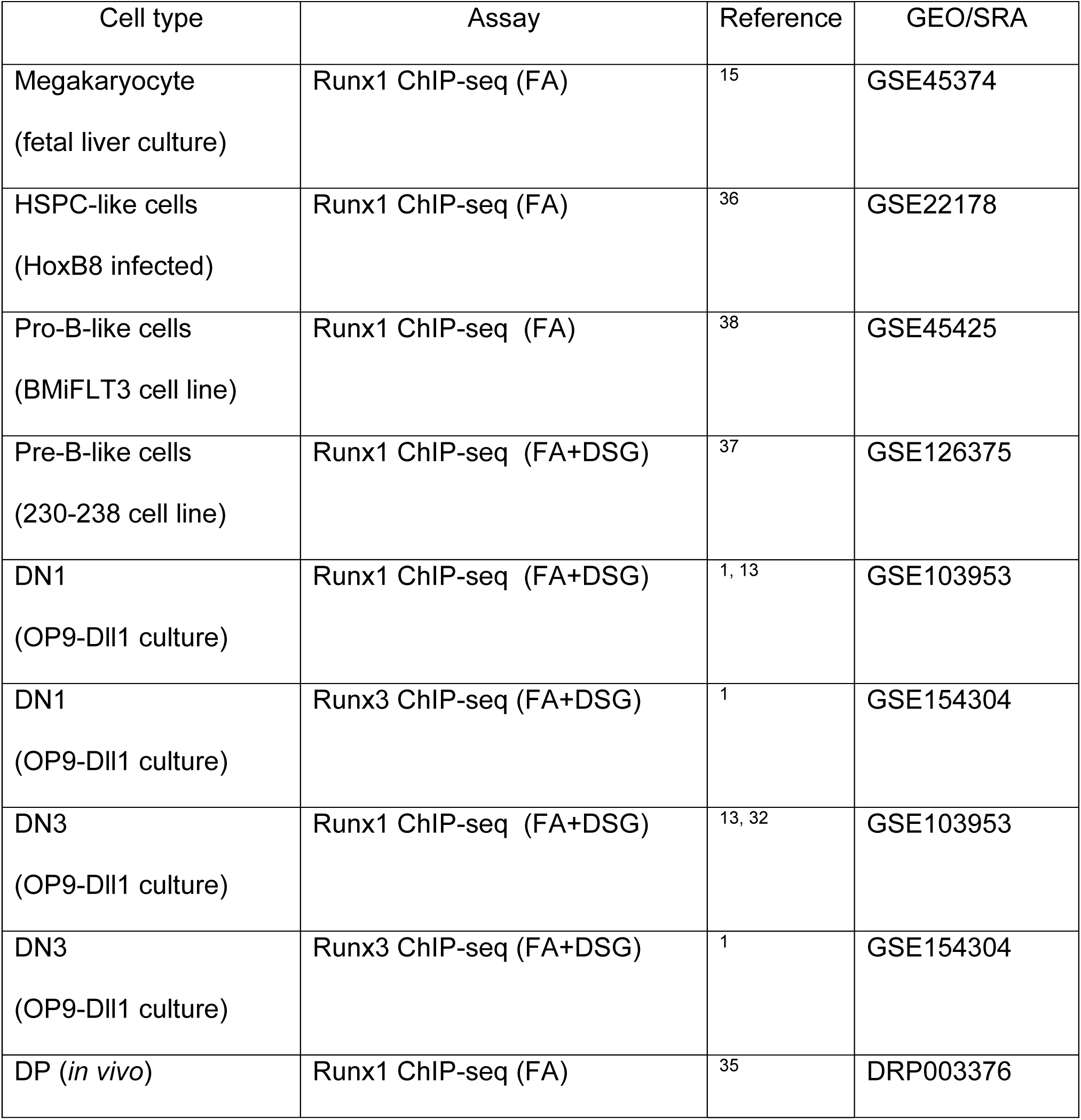
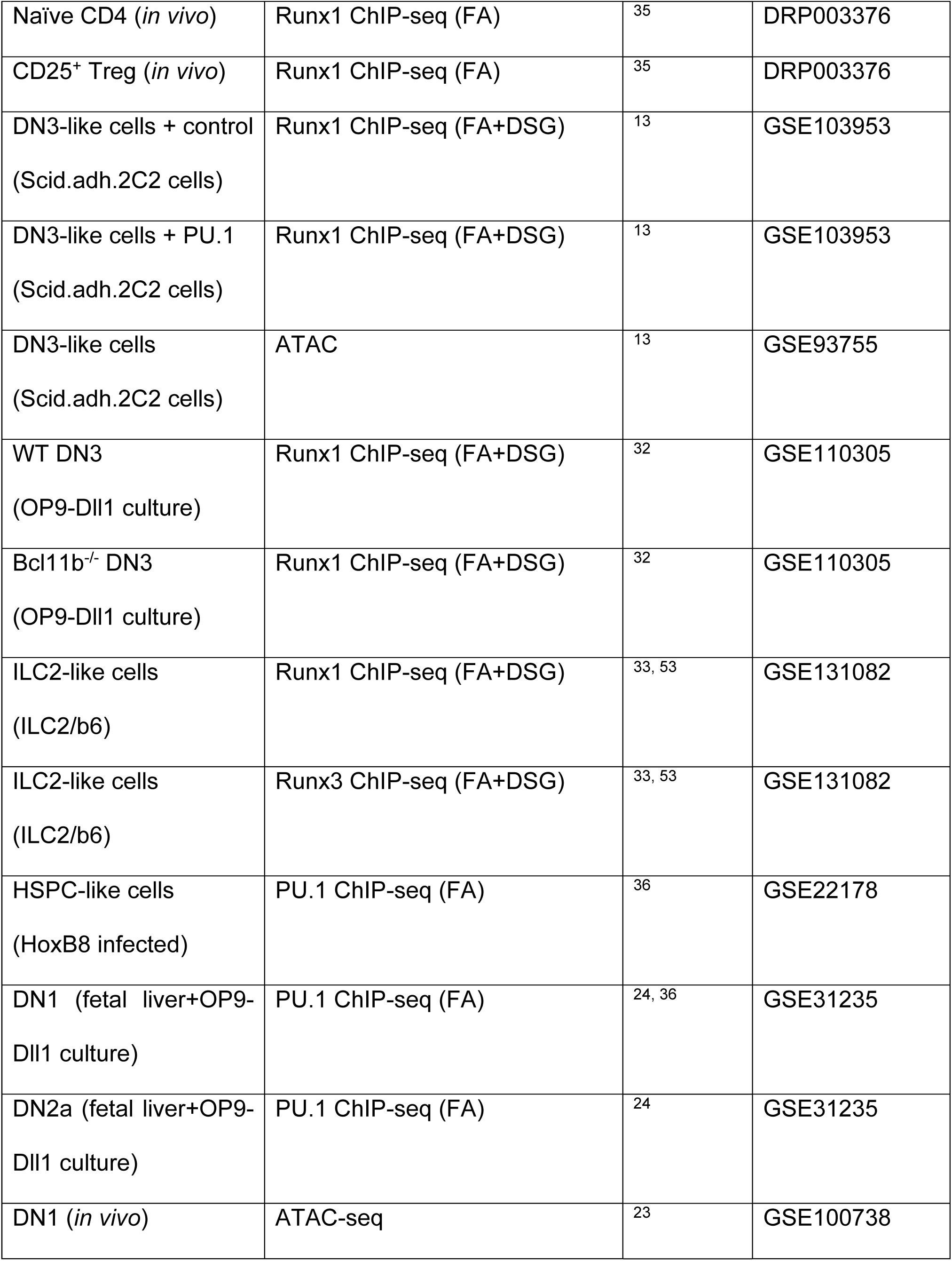
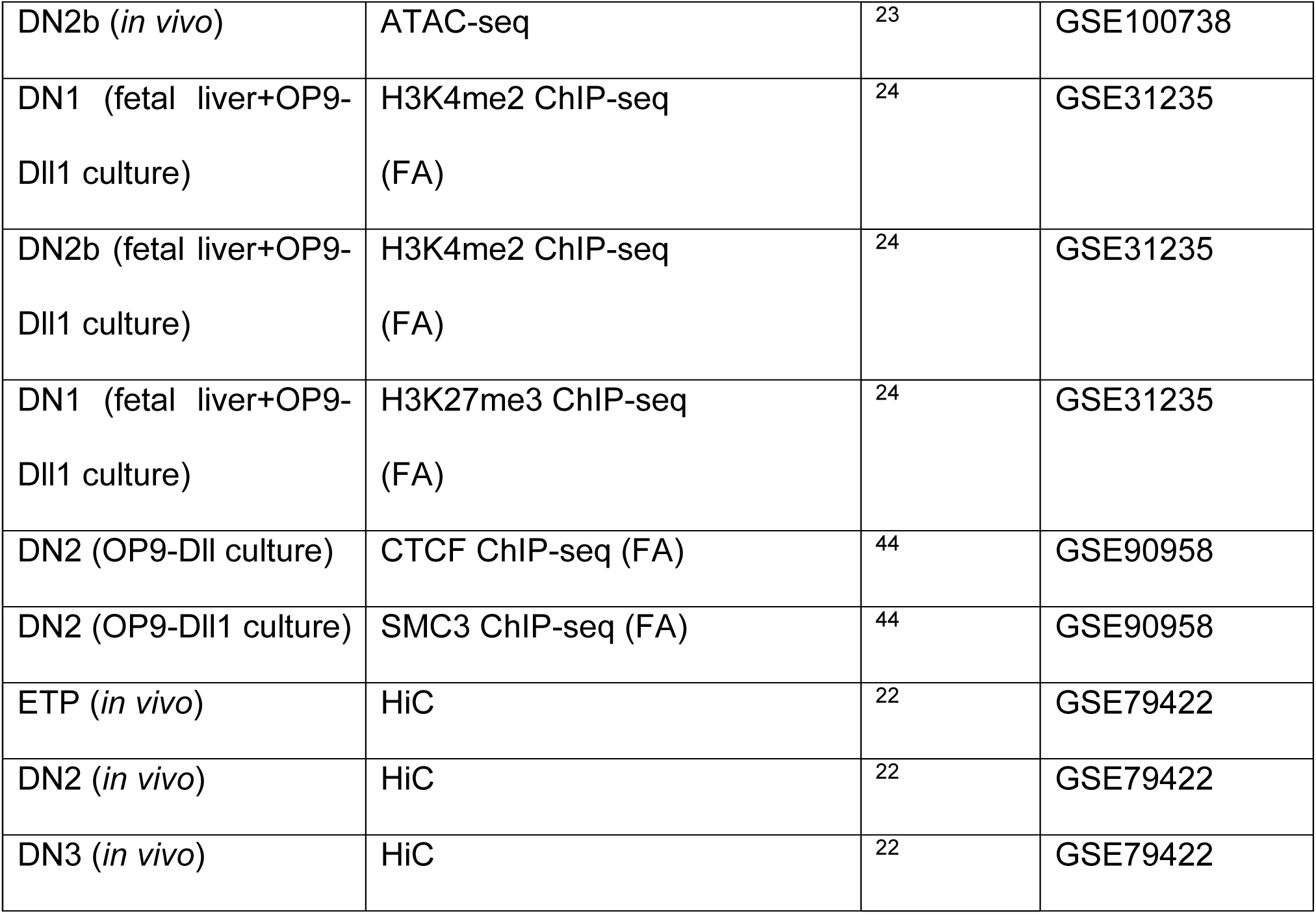

| Cell type | Assay | Reference | GEO/SRA |
| --- | --- | --- | --- |
| Megakaryocyte<br>(fetal liver culture) | Runx1 ChIP-seq (FA) | <sup>15</sup> | GSE45374 |
| HSPC-like cells<br>(HoxB8 infected) | Runx1 ChIP-seq (FA) | <sup>36</sup> | GSE22178 |
| Pro-B-like cells<br>(BMiFLT3 cell line) | Runx1 ChIP-seq (FA) | <sup>38</sup> | GSE45425 |
| Pre-B-like cells<br>(230-238 cell line) | Runx1 ChIP-seq (FA+DSG) | <sup>37</sup> | GSE126375 |
| DN1<br>(OP9-DII1 culture) | Runx1 ChIP-seq (FA+DSG) | <sup>1, 13</sup> | GSE103953 |
| DN1<br>(OP9-DII1 culture) | Runx3 ChIP-seq (FA+DSG) | <sup>1</sup> | GSE154304 |
| DN3<br>(OP9-DII1 culture) | Runx1 ChIP-seq (FA+DSG) | <sup>13, 32</sup> | GSE103953 |
| DN3<br>(OP9-DII1 culture) | Runx3 ChIP-seq (FA+DSG) | <sup>1</sup> | GSE154304 |
| DP (*in vivo*) | Runx1 ChIP-seq (FA) | <sup>35</sup> | DRP003376 |
| Naïve CD4 (*in vivo*) | Runx1 ChIP-seq (FA) | 35 | DRP003376 |
| CD25<sup>+</sup> Treg (*in vivo*) | Runx1 ChIP-seq (FA) | 35 | DRP003376 |
| DN3-like cells + control<br>(Scid.adh.2C2 cells) | Runx1 ChIP-seq (FA+DSG) | 13 | GSE103953 |
| DN3-like cells + PU.1<br>(Scid.adh.2C2 cells) | Runx1 ChIP-seq (FA+DSG) | 13 | GSE103953 |
| DN3-like cells<br>(Scid.adh.2C2 cells) | ATAC | 13 | GSE93755 |
| WT DN3<br>(OP9-Dll1 culture) | Runx1 ChIP-seq (FA+DSG) | 32 | GSE110305 |
| Bcl11b<sup>-/-</sup> DN3<br>(OP9-Dll1 culture) | Runx1 ChIP-seq (FA+DSG) | 32 | GSE110305 |
| ILC2-like cells<br>(ILC2/b6) | Runx1 ChIP-seq (FA+DSG) | 33, 53 | GSE131082 |
| ILC2-like cells<br>(ILC2/b6) | Runx3 ChIP-seq (FA+DSG) | 33, 53 | GSE131082 |
| HSPC-like cells<br>(HoxB8 infected) | PU.1 ChIP-seq (FA) | 36 | GSE22178 |
| DN1 (fetal liver+OP9-<br>Dll1 culture) | PU.1 ChIP-seq (FA) | 24, 36 | GSE31235 |
| DN2a (fetal liver+OP9-<br>Dll1 culture) | PU.1 ChIP-seq (FA) | 24 | GSE31235 |
| DN1 (*in vivo*) | ATAC-seq | 23 | GSE100738 |
| DN2b (*in vivo*) | ATAC-seq | <sup>23</sup> | GSE100738 |
| DN1 (fetal liver+OP9-Dll1 culture) | H3K4me2 ChIP-seq (FA) | <sup>24</sup> | GSE31235 |
| DN2b (fetal liver+OP9-Dll1 culture) | H3K4me2 ChIP-seq (FA) | <sup>24</sup> | GSE31235 |
| DN1 (fetal liver+OP9-Dll1 culture) | H3K27me3 ChIP-seq (FA) | <sup>24</sup> | GSE31235 |
| DN2 (OP9-Dll culture) | CTCF ChIP-seq (FA) | <sup>44</sup> | GSE90958 |
| DN2 (OP9-Dll1 culture) | SMC3 ChIP-seq (FA) | <sup>44</sup> | GSE90958 |
| ETP (*in vivo*) | HiC | <sup>22</sup> | GSE79422 |
| DN2 (*in vivo*) | HiC | <sup>22</sup> | GSE79422 |
| DN3 (*in vivo*) | HiC | <sup>22</sup> | GSE79422 |

### Note on CUT&RUN comparison with DSG-assisted ChIP-seq

C&R^39, 40, 78^ was used throughout this study to track Runx binding, not only because of its tolerance for low cell numbers but also because the ability to omit crosslinking should focus the analysis on sites of direct Runx-DNA binding. Fig. S1 shows control analyses in which we directly compared Runx binding profiles from unperturbed DN1 (Phase 1) and DN2b/DN3 (Phase 2) cells using C&R or using ChIP-seq (disuccinimidyl glutarate (DSG) + formaldehyde cross-linked for better efficiency^1^) (Fig. S1c). A large number of Runx binding sites detected by C&R and ChIP-seq agreed, but C&R detected a smaller number of occupancies than the ChIP-seq, especially detecting fewer promoter regions. In addition, the non-promoter Runx occupancies detected by ChIP-seq but not by C&R had globally weaker Runx motifs. This supports our prediction that indirect binding would be better captured by ChIP-seq due to protein-protein as well as protein-DNA cross-links. In contrast, the Runx occupancy sites more efficiently detected by C&R displayed high quality Runx motifs with less frequent ETS motif co-enrichment than the ChIP-seq preferentially detected sites (Fig. S1d, e). Finally, C&R efficiently detected Runx occupancies in both open and closed chromatin sites (Fig. S1f). Thus, direct Runx binding sites, especially in non-promoter regions, could be reliably determined by C&R.

### ChromHMM for analysis of local chromatin states and long-range analysis of chromatin A/B compartments

The chromatin states of pro-T cells were inferred utilizing ChromHMM (v. 1.23)^42, 43^ by learning models using previously published histone mark ChIP-seq data^24^ and CTCF, SMC3 ChIP-seq^44^.ChromHMM calculates the most probable state for each genomic segment based on a multivariate hidden Markov model (HMM)^42, 43^. For our ChromHMM analysis, we utilized ATAC-seq (chromatin accessibility), H3K4me2 marks (active histone), and H3K27me3 marks (repressive histone), along with CTCF and SMC3 (DNA loop-forming machineries) binding data obtained from DN1 (representing pre-commitment stage) and DN2b cells (representing post-commitment stage)^23, 24, 44^. ChromHMM defined 20 different chromatin states in pro-T cells, which included Phase 1-preferential active sites (state 1-4), Phase 2-preferential active sites (state 5-7), active sites in both stages (state 8-11, 14), weakly repressed or bivalent regions (state 12, 13), and the sites that were repressed in all stages (state 15, 16) (Fig. S2c). For computing chromatin state using ChromHMM, we followed the authors’ instructions^42, 43^. Briefly, replicated DNA sequencing bam files were merged using Samtools, then binarized using a ChromHMM function (BinarizeBam). Using binarized bam files, the chromatin state models were calculated for the mm10 genome (LearnModel). To compare association between the computed chromatin states and different groups of Runx binding sites, previously defined Group 1, 2, 3, and promoter Runx binding sites were provided as a set of external annotation data, and the enrichment was calculated (OverlapEnrichment).

The A/B compartment analysis was performed by using previously reported HiC data^22^ after converting mm9 20kb-bin tracks (GSE79422) to mm9 1kb-bin tracks, and then lifting over mm9 1kb-bin tracks to mm10 1kb-bin tracks.

### Single cell RNA-seq analyses

The raw reads from cDNA and hashtag oligo libraries were processed as previously described^55^. Briefly, cDNA libraries were aligned to the mouse reference genome GRCm38/mm10 using CellRanger3 and the hashtag oligo libraries were quantified and demultiplexed using in-house tools (hashtag_tool)^55^. Seurat v4^85^ was utilized for QC and downstream analysis. For QC, singlets (cells displaying unique hashtag oligo identity) expressing at least 1300 genes (transcript > 1) were considered. Outlier cells expressing more than 6800 genes (potential doublets) and displaying high mitochondrial RNA contents (> 1 %) were further filtered. Two independent experiments were integrated with reciprocal principal component analysis (PCA) algorithm using the 5000 anchor features. After data integration, PC analysis was performed, and the first 30 PCs were utilized for computing tSNE and UMAP parameters. The pathway enrichment in each cell was conducted with single-sample gene set enrichment (ssGSEA) tool (https://github.com/GSEA-MSigDB/ssGSEA-gpmodule). The pseudotime inference was performed using Monocle3 ^86, 87^ by defining the root principal node based on the known gene expression pattern of early thymic progenitors (Flt3>5 & Kit>1 & Lmo2 > 1 & Il2ra<0.1 & Tcf7<0.1). To determine the UMAP2 acceleration time window, first, linear regression was performed using UMAP2 values and pseudotime scores from control cells. Then, residual values (predicted pseudotime score based on linear regression fit vs. the observed pseudotime score) were calculated. UMAP2 window -30 to 5 was chosen as more than 50% of residuals of OE cells were greater than interquartile residual values. For visualization of Seurat4 and Monocle3 analysis results, ggplot2 (v. 3.3.5) and cowplot (v. 1.1.1) packages were utilized.

To infer gene regulatory network (GRN), integrated Seurat object was converted to loom files using SeuratDisk and SeuratData, and then pySCENIC (v.0.11.2)^60, 61^ was employed to compute co-expression network and search for potential direct target genes. The default parameters and the standard workflows were applied. Results were visualized with matplotlib (v.3.5.3).

### Differentially Expressed Gene analysis

To define differentially expressed genes (DEGs), we first excluded alternative lineage clusters (clusters 12-16) to focus on the cells on the T-developmental pathway. Differential expression tests were conducted using Wilcox test employing a Seurat tool (FindMarkers) with pseudocount = 0.1, min.pct=0.2, min.cells.group=3, min.cells.feature=3 parameters. Genes displaying absolute fold-change > 1.5, adjusted p-value<0.001 were considered as DEGs. “Expressed” non-DEGs were defined by 1) their detection via scRNA-seq and then 2) not sensitive to any of Runx perturbation (KO nor OE) at any timepoints. Runx core-DEGs were defined if a given gene is sensitive to both Runx1 OE and Runx1/Runx3 KO perturbations for activation or inhibition. See Table S1 and S2 for the full lists of DEGs from this study and the comparison with previously published bulk-RNAseq results. Note that bulk-RNAseq results do not exclude cells consisting alternative lineage clusters.

### C&R and Runx target gene association analysis

To test whether a gain or loss of Runx binding is associated with Runx-mediated gene regulation, different groups of Runx peaks were annotated with putatively associated genes using GREAT and enrichment patterns were calculated as described previously^1^. Briefly, presence of any non-promoter Runx peak(s) in surrounding genomic regions of each transcript in DEG and non-DEG groups was scored. Then, the following statistics were further examined: (1) the percentage of genes in each category linked to Runx binding, (2) whether Runx binding is equally or not equally distributed between DEG vs. non-DEGs by performing Fisher’s exact test, (3) whether different classes of DEGs (activated or inhibited, scored by KO or OE or both) had preferential enrichment for a certain group of Runx binding using the z-score, which is calculated by standardized residual analysis (Fig. S7a). To calculated composition of different groups of Runx binding, the number of peaks in a given category was divided by the total number of Runx peaks annotated to each gene, and the percent of each group of peaks was reported (Fig. S7b).

### Other statistical tests

Nonparametric tests comparing two distributions were performed by two-sample Kolmogorov-Smirnov test (Fig.1f, 6d, 8d, S1d, S3e). To compare the average of two groups, t-test was performed (Fig. 2d)). To compare the average of three or more groups, One-way ANOVA with Tukey’s multiple comparisons test was used (Fig. S3c). To test the effect of two independent variables on a dependent variable, Two-way ANOVA was utilized (Fig. 2b, 2e, 5b, 5e, 5f, 7e, S3b, S3f, S5b, S5c). The Kruskal–Wallis test by ranks was used to compare pseudotime progression rate (Fig. 4b). To assess linear correlation between two different parameters, Pearson correlation coefficient (Pearson’s *r*) was calculated (Fig. 4c). The monotonic correlation between two parameters were tested using Spearman’s rank correlation coefficient (Fig. 4c). Linear regression analysis was conducted to find the best fit line for UMAP-2 values and Pseudotime scores of Control groups (Fig. 4c). Fisher’s exact test and standardized residual analysis was conducted to evaluate association between categorical variables (Fig, S4c, S7a).

One-way ANOVA, Two-way ANOVA, t-test, and the Kruskal-Wallis test were performed using Prism software (v.9.4.1, GraphPad). Two-sample Kolmogorov-Smirnov test, Pearson’s *r* calculation, and Spearman’s rank ρ calculation were performed using scipy.stats. (v.1.9.1) from Python (v.3.8.13). Linear regression analysis, Fisher’s exact test, and standardized residual analysis were performed using R (v.4.1.1). *p < 0.05; **p < 0.01; ***p <0.001 for t-test, One-way ANOVA, Two-way ANOVA, and Kolmogorov-Smirnov test. * |z-score| > 1.9599; ** |z-score| > 2.5758; *** |z-score| > 3.2905 for standardized residual analysis.

## References

1. Shin, B., Hosokawa, H., Romero-Wolf, M., Zhou, W., Masuhara, K., Tobin, V.R., Levanon, D., Groner, Y. & Rothenberg, E.V. Runx1 and Runx3 drive progenitor to T-lineage transcriptome conversion in mouse T cell commitment via dynamic genomic site switching. Proc Natl Acad Sci U S A 118 (2021).

2. Guo, Y., Maillard, I., Chakraborti, S., Rothenberg, E.V. & Speck, N.A. Core binding factors are necessary for natural killer cell development and cooperate with Notch signaling during T-cell specification. Blood 112, 480–92 (2008).

3. Zhao, L., Cannons, J.L., Anderson, S., Kirby, M., Xu, L., Castilla, L.H., Schwartzberg, P.L., Bosselut, R. & Liu, P.P. CBFB-MYH11 hinders early T-cell development and induces massive cell death in the thymus. Blood 109, 3432–3440 (2007).

4. Taniuchi, I., Osato, M., Egawa, T., Sunshine, M.J., Bae, S.C., Komori, T., Ito, Y. & Littman, D.R. Differential requirements for Runx proteins in CD4 repression and epigenetic silencing during T lymphocyte development. Cell 111, 621–33 (2002).

5. Egawa, T., Tillman, R.E., Naoe, Y., Taniuchi, I. & Littman, D.R. The role of the Runx transcription factors in thymocyte differentiation and in homeostasis of naive T cells. J Exp Med 204, 1945–57 (2007).

6. Oestreich, K.J., Cobb, R.M., Pierce, S., Chen, J., Ferrier, P. & Oltz, E.M. Regulation of TCRbeta gene assembly by a promoter/enhancer holocomplex. Immunity 24, 381–91 (2006).

7. Yzaguirre, A.D., de Bruijn, M.F. & Speck, N.A. The Role of Runx1 in Embryonic Blood Cell Formation. Adv Exp Med Biol 962, 47–64 (2017).

8. Growney, J.D., Shigematsu, H., Li, Z., Lee, B.H., Adelsperger, J., Rowan, R., Curley, D.P., Kutok, J.L., Akashi, K., Williams, I.R. et al. Loss of Runx1 perturbs adult hematopoiesis and is associated with a myeloproliferative phenotype. Blood 106, 494–504 (2005).

9. Ichikawa, M., Asai, T., Saito, T., Seo, S., Yamazaki, I., Yamagata, T., Mitani, K., Chiba, S., Ogawa, S., Kurokawa, M. & Hirai, H. AML-1 is required for megakaryocytic maturation and lymphocytic differentiation, but not for maintenance of hematopoietic stem cells in adult hematopoiesis. Nat Med 10, 299–304 (2004).

10. Talebian, L., Li, Z., Guo, Y., Gaudet, J., Speck, M.E., Sugiyama, D., Kaur, P., Pear, W.S., Maillard, I. & Speck, N.A. T-lymphoid, megakaryocyte, and granulocyte development are sensitive to decreases in CBFb dosage. Blood 109, 11–21 (2007).

11. Seo, W., Ikawa, T., Kawamoto, H. & Taniuchi, I. Runx1-Cbfβ facilitates early B lymphocyte development by regulating expression of Ebf1. J Exp Med 209, 1255–62 (2012).

12. Pham, T.-H., Minderjahn, J., Schmidl, C., Hoffmeister, H., Schmidhofer, S., Chen, W., Langst, G., Benner, C. &Rehli, M. Mechanisms of in vivo binding site selection of the hematopoietic master transcription factor PU.1. Nucleic Acids Res 41, 6391–6402 (2013).

13. Ungerback, J., Hosokawa, H., Wang, X., Strid, T., Williams, B.A., Sigvardsson, M. & Rothenberg, E.V. Pioneering, chromatin remodeling, and epigenetic constraint in early T-cell gene regulation by SPI1 (PU.1). Genome Res 28, 1508–1519 (2018).

14. Lin, Q., Chauvistré, H., Costa, I.G., Gusmao, E.G., Mitzka, S., Hanzelmann, S., Baying, B., Klisch, T., Moriggl, R., Hennuy, B. et al. Epigenetic program and transcription factor circuitry of dendritic cell development. Nucleic Acids Res 43, 9680–93 (2015).

15. Pencovich, N., Jaschek, R., Dicken, J., Amit, A., Lotem, J., Tanay, A. & Groner, Y. Cell-autonomous function of Runx1 transcriptionally regulates mouse megakaryocytic maturation. PLoS One 8, e64248 (2013).

16. Zang, C., Luyten, A., Chen, J., Liu, X.S. & Shivdasani, R.A. NF-E2, FLI1 and RUNX1 collaborate at areas of dynamic chromatin to activate transcription in mature mouse megakaryocytes. Sci Rep 6, 30255 (2016).

17. Miyazaki, M., Rivera, R.R., Miyazaki, K., Lin, Y.C., Agata, Y. & Murre, C. The opposing roles of the transcription factor E2A and its antagonist Id3 that orchestrate and enforce the naive fate of T cells. Nat Immunol 12, 992–1001 (2011).

18. Lin, Y.C., Jhunjhunwala, S., Benner, C., Heinz, S., Welinder, E., Mansson, R., Sigvardsson, M., Hagman, J., Espinoza, C.A., Dutkowski, J. et al. A global network of transcription factors, involving E2A, EBF1 and Foxo1, that orchestrates B cell fate. Nat.Immunol. 11, 635–643 (2010).

19. Zhong, Y., Walker, S.K., Pritykin, Y., Leslie, C.S., Rudensky, A.Y. & van der Veeken, J. Hierarchical regulation of the resting and activated T cell epigenome by major transcription factor families. Nat Immunol 23, 122–134 (2022).

20. Yui, M.A. & Rothenberg, E.V. Developmental gene networks: a triathlon on the course to T cell identity. Nat Rev Immunol 14, 529–545 (2014).

21. Hosokawa, H. & Rothenberg, E.V. How transcription factors drive choice of the T cell fate. Nat Rev Immunol 21, 162–176 (2021).

22. Hu, G., Cui, K., Fang, D., Hirose, S., Wang, X., Wangsa, D., Jin, W., Ried, T., Liu, P., Zhu, J. et al. Transformation of Accessible Chromatin and 3D Nucleome Underlies Lineage Commitment of Early T Cells. Immunity 48, 227–242 e8 (2018).

23. Yoshida, H., Lareau, C.A., Ramirez, R.N., Rose, S.A., Maier, B., Wroblewska, A., Desland, F., Chudnovskiy, A., Mortha, A., Dominguez, C. et al. The cis-Regulatory Atlas of the Mouse Immune System. Cell 176, 897–912 e20 (2019).

24. Zhang, J.A., Mortazavi, A., Williams, B.A., Wold, B.J. & Rothenberg, E.V. Dynamic transformations of genome-wide epigenetic marking and transcriptional control establish T cell identity. Cell 149, 467–82 (2012).

25. Zhou, W., Yui, M.A., Williams, B.A., Yun, J., Wold, B.J., Cai, L. & Rothenberg, E.V. Single-Cell Analysis Reveals Regulatory Gene Expression Dynamics Leading to Lineage Commitment in Early T Cell Development. Cell Syst 9, 321–337 e9 (2019).

26. Mingueneau, M., Kreslavsky, T., Gray, D., Heng, T., Cruse, R., Ericson, J., Bendall, S., Spitzer, M.H., Nolan, G.P., Kobayashi, K. et al. The transcriptional landscape of αβ T cell differentiation. Nat Immunol 14, 619–632 (2013).

27. Bruno, L., Ramlall, V., Studer, R.A., Sauer, S., Bradley, D., Dharmalingam, G., Carroll, T., Ghoneim, M., Chopin, M., Nutt, S.L. et al. Selective deployment of transcription factor paralogs with submaximal strength facilitates gene regulation in the immune system. Nat Immunol 20, 1372–1380 (2019).

28. Longabaugh, W.J.R., Zeng, W., Zhang, J.A., Hosokawa, H., Jansen, C.S., Li, L., Romero-Wolf, M., Liu, P., Kueh, H.Y., Mortazavi, A. & Rothenberg, E.V. Bcl11b and combinatorial resolution of cell fate in the T-cell gene regulatory network. Proc Natl Acad Sci U S A 114, 5800–5807 (2017).

29. Rothenberg, E.V. Programming for T-lymphocyte fates: modularity and mechanisms. Genes Dev 33, 1117–1135 (2019).

30. Rothenberg, E.V. Logic and lineage impacts on functional transcription factor deployment for T-cell fate commitment. Biophys J 120, 4162–4181 (2021).

31. Naito, T., Tanaka, H., Naoe, Y. & Taniuchi, I. Transcriptional control of T-cell development. Int Immunol 23, 661–8 (2011).

32. Hosokawa, H., Romero-Wolf, M., Yui, M.A., Ungerback, J., Quiloan, M.L.G., Matsumoto, M., Nakayama, K.I., Tanaka, T. & Rothenberg, E.V. Bcl11b sets pro-T cell fate by site-specific cofactor recruitment and by repressing Id2 and Zbtb16. Nat Immunol 19, 1427–1440 (2018).

33. Hosokawa, H., Koizumi, M., Masuhara, K., Romero-Wolf, M., Tanaka, T., Nakayama, T. & Rothenberg, E.V. Stage-specific action of Runx1 and GATA3 controls silencing of PU.1 expression in mouse pro-T cells. J Exp Med 218 (2021).

34. Hamey, F.K., Nestorowa, S., Kinston, S.J., Kent, D.G., Wilson, N.K. & Gottgens, B. Reconstructing blood stem cell regulatory network models from single-cell molecular profiles. Proc Natl Acad Sci U S A 114, 5822–5829 (2017).

35. Kitagawa, Y., Ohkura, N., Kidani, Y., Vandenbon, A., Hirota, K., Kawakami, R., Yasuda, K., Motooka, D., Nakamura, S., Kondo, M. et al. Guidance of regulatory T cell development by Satb1-dependent super-enhancer establishment. Nat Immunol 18, 173–183 (2017).

36. Wilson, N.K., Foster, S.D., Wang, X., Knezevic, K., Schutte, J., Kaimakis, P., Chilarska, P.M., Kinston, S., Ouwehand, W.H., Dzierzak, E. et al. Combinatorial transcriptional control in blood stem/progenitor cells: genome-wide analysis of ten major transcriptional regulators. Cell Stem Cell 7, 532–44 (2010).

37. Okuyama, K., Strid, T., Kuruvilla, J., Somasundaram, R., Cristobal, S., Smith, E., Prasad, M., Fioretos, T., Lilljebjorn, H., Soneji, S. et al. PAX5 is part of a functional transcription factor network targeted in lymphoid leukemia. PLoS Genet 15, e1008280 (2019).

38. Niebuhr, B., Kriebitzsch, N., Fischer, M., Behrens, K., Gunther, T., Alawi, M., Bergholz, U., Muller, U., Roscher, S., Ziegler, M. et al. Runx1 is essential at two stages of early murine B-cell development. Blood 122, 413–23 (2013).

39. Skene, P.J., Henikoff, J.G. & Henikoff, S. Targeted in situ genome-wide profiling with high efficiency for low cell numbers. Nat Protoc 13, 1006–1019 (2018).

40. Skene, P.J. & Henikoff, S. An efficient targeted nuclease strategy for high-resolution mapping of DNA binding sites. Elife 6 (2017).

41. Lieberman-Aiden, E., van Berkum, N.L., Williams, L., Imakaev, M., Ragoczy, T., Telling, A., Amit, I., Lajoie, B.R., Sabo, P.J., Dorschner, M.O. et al. Comprehensive mapping of long-range interactions reveals folding principles of the human genome. Science 326, 289–93 (2009).

42. Ernst, J. & Kellis, M. Chromatin-state discovery and genome annotation with ChromHMM. Nat Protoc 12, 2478–2492 (2017).

43. Ernst, J. & Kellis, M. ChromHMM: automating chromatin-state discovery and characterization. Nat Methods 9, 215–6 (2012).

44. Isoda, T., Moore, A.J., He, Z., Chandra, V., Aida, M., Denholtz, M., Piet van Hamburg, J., Fisch, K.M., Chang, A.N., Fahl, S.P., et al. Non-coding Transcription Instructs Chromatin Folding and Compartmentalization to Dictate Enhancer-Promoter Communication and T Cell Fate. Cell 171, 103–119 e18 (2017).

45. Bonifer, C., Levantini, E., Kouskoff, V. & Lacaud, G. Runx1 Structure and Function in Blood Cell Development. Adv Exp Med Biol 962, 65–81 (2017).

46. Friedman, A.D. Cell cycle and developmental control of hematopoiesis by Runx1. J Cell Physiol 219, 520–524 (2009).

47. Yu, M., Mazor, T., Huang, H., Huang, H.T., Kathrein, K.L., Woo, A.J., Chouinard, C.R., Labadorf, A., Akie, T.E., Moran, T.B. et al. Direct recruitment of polycomb repressive complex 1 to chromatin by core binding transcription factors. Mol Cell 45, 330–43 (2012).

48. Schmitt, T.M. & Zúñiga-Pflücker, J.C. Induction of T cell development from hematopoietic progenitor cells by Delta-like-1 in vitro. Immunity 17, 749–56 (2002).

49. Kueh, H.Y., Yui, M.A., Ng, K.K., Pease, S.S., Zhang, J.A., Damle, S.S., Freedman, G., Siu, S., Bernstein, I.D., Elowitz, M.B. & Rothenberg, E.V. Asynchronous combinatorial action of four regulatory factors activates Bcl11b for T cell commitment. Nat Immunol 17, 956–65 (2016).

50. Wong, W.F., Nakazato, M., Watanabe, T., Kohu, K., Ogata, T., Yoshida, N., Sotomaru, Y., Ito, M., Araki, K., Telfer, J. et al. Over-expression of Runx1 transcription factor impairs the development of thymocytes from the double-negative to double-positive stages. Immunology 130, 243–53 (2010).

51. Sambandam, A., Maillard, I., Zediak, V.P., Xu, L., Gerstein, R.M., Aster, J.C., Pear, W.S. & Bhandoola, A. Notch signaling controls the generation and differentiation of early T lineage progenitors. Nat Immunol 6, 663–70 (2005).

52. Ramond, C., Berthault, C., Burlen-Defranoux, O., de Sousa, A.P., Guy-Grand, D., Vieira, P., Pereira, P. & Cumano, A. Two waves of distinct hematopoietic progenitor cells colonize the fetal thymus. Nat Immunol 15, 27–35 (2014).

53. Hosokawa, H., Romero-Wolf, M., Yang, Q., Motomura, Y., Levanon, D., Groner, Y., Moro, K., Tanaka, T. & Rothenberg, E.V. Cell type-specific actions of Bcl11b in early T-lineage and group 2 innate lymphoid cells. J Exp Med 217, e20190972 (2020).

54. Miyazaki, M., Miyazaki, K., Chen, K., Jin, Y., Turner, J., Moore, A.J., Saito, R., Yoshida, K., Ogawa, S., Rodewald, H.R. et al. The E-Id Protein Axis Specifies Adaptive Lymphoid Cell Identity and Suppresses Thymic Innate Lymphoid Cell Development. Immunity 46, 818–834 e4 (2017).

55. Zhou, W., Gao, F., Romero-Wolf, M., Jo, S. & Rothenberg, E.V. Single-cell deletion analyses show control of pro-T cell developmental speed and pathways by Tcf7, Spi1, Gata3, Bcl11a, Erg, and Bcl11b. Sci Immunol 7, eabm1920 (2022).

56. Wiede, F., Dudakov, J.A., Lu, K.H., Dodd, G.T., Butt, T., Godfrey, D.I., Strasser, A., Boyd, R.L. & Tiganis, T. PTPN2 regulates T cell lineage commitment and αβ versus γδ specification. J Exp Med 214, 2733–2758 (2017).

57. Yao, Z., Cui, Y., Watford, W.T., Bream, J.H., Yamaoka, K., Hissong, B.D., Li, D., Durum, S.K., Jiang, Q., Bhandoola, A. et al. Stat5a/b are essential for normal lymphoid development and differentiation. Proc Natl Acad Sci U S A 103, 1000–5 (2006).

58. Seet, C.S., He, C., Bethune, M.T., Li, S., Chick, B., Gschweng, E.H., Zhu, Y., Kim, K., Kohn, D.B., Baltimore, D. et al. Generation of mature T cells from human hematopoietic stem and progenitor cells in artificial thymic organoids. Nat Methods 14, 521–530 (2017).

59. Montel-Hagen, A., Sun, V., Casero, D., Tsai, S., Zampieri, A., Jackson, N., Li, S., Lopez, S., Zhu, Y., Chick, B. et al. In Vitro Recapitulation of Murine Thymopoiesis from Single Hematopoietic Stem Cells. Cell Rep 33, 108320 (2020).

60. Aibar, S., Gonzalez-Blas, C.B., Moerman, T., Huynh-Thu, V.A., Imrichova, H., Hulselmans, G., Rambow, F., Marine, J.C., Geurts, P., Aerts, J. et al. SCENIC: single-cell regulatory network inference and clustering. Nat Methods 14, 1083–1086 (2017).

61. Van de Sande, B., Flerin, C., Davie, K., De Waegeneer, M., Hulselmans, G., Aibar, S., Seurinck, R., Saelens, W., Cannoodt, R., Rouchon, Q. et al. A scalable SCENIC workflow for single-cell gene regulatory network analysis. Nat Protoc 15, 2247–2276 (2020).

62. Wunderlich, Z. & Mirny, L.A. Different gene regulation strategies revealed by analysis of binding motifs. Trends Genet 25, 434–40 (2009).

63. Kribelbauer, J.F., Rastogi, C., Bussemaker, H.J. & Mann, R.S. Low-Affinity Binding Sites and the Transcription Factor Specificity Paradox in Eukaryotes. Annu Rev Cell Dev Biol 35, 357–379 (2019).

64. Crocker, J., Abe, N., Rinaldi, L., McGregor, A.P., Frankel, N., Wang, S., Alsawadi, A., Valenti, P., Plaza, S., Payre, F. et al. Low affinity binding site clusters confer hox specificity and regulatory robustness. Cell 160, 191–203 (2015).

65. Farley, E.K., Olson, K.M., Zhang, W., Brandt, A.J., Rokhsar, D.S. & Levine, M.S. Suboptimization of developmental enhancers. Science 350, 325–8 (2015).

66. Boisclair Lachance, J.F., Webber, J.L., Hong, L., Dinner, A.R. & Rebay, I. Cooperative recruitment of Yan via a high-affinity ETS supersite organizes repression to confer specificity and robustness to cardiac cell fate specification. Genes Dev 32, 389–401 (2018).

67. Gao, A., Shrinivas, K., Lepeudry, P., Suzuki, H.I., Sharp, P.A. & Chakraborty, A.K. Evolution of weak cooperative interactions for biological specificity. Proc Natl Acad Sci U S A 115, E11053–E11060 (2018).

68. Brewster, R.C., Weinert, F.M., Garcia, H.G., Song, D., Rydenfelt, M. & Phillips, R. The transcription factor titration effect dictates level of gene expression. Cell 156, 1312–1323 (2014).

69. Lie-A-Ling, M., Marinopoulou, E., Lilly, A.J., Challinor, M., Patel, R., Lancrin, C., Kouskoff, V. & Lacaud, G. Regulation of RUNX1 dosage is crucial for efficient blood formation from hemogenic endothelium. Development 145 (2018).

70. Lacaud, G., Kouskoff, V., Trumble, A., Schwantz, S. & Keller, G. Haploinsufficiency of *Runx1* results in the acceleration of mesodermal development and hemangioblast specification upon in vitro differentiation of ES cells. Blood 103, 886–889 (2004).

71. Telfer, J.C., Hedblom, E.E., Anderson, M.K., Laurent, M.N. & Rothenberg, E.V. Localization of the domains in Runx transcription factors required for the repression of CD4 in thymocytes. J Immunol 172, 4359–70 (2004).

72. Romero-Wolf, M., Shin, B., Zhou, W., Koizumi, M., Rothenberg, E.V. & Hosokawa, H. Notch2 complements Notch1 to mediate inductive signaling that initiates early T cell development. J Cell Biol 219, e202005093 (2020).

73. Ng, K.K., Yui, M.A., Mehta, A., Siu, S., Irwin, B., Pease, S., Hirose, S., Elowitz, M.B., Rothenberg, E.V. & Kueh, H.Y. A stochastic epigenetic switch controls the dynamics of T-cell lineage commitment. Elife 7 (2018).

74. Dionne, C.J., Tse, K.Y., Weiss, A.H., Franco, C.B., Wiest, D.L., Anderson, M.K. & Rothenberg, E.V. Subversion of T lineage commitment by PU.1 in a clonal cell line system. Dev Biol 280, 448–66 (2005).

75. Franco, C.B., Scripture-Adams, D.D., Proekt, I., Taghon, T., Weiss, A.H., Yui, M.A., Adams, S.L., Diamond, R.A. & Rothenberg, E.V. Notch/Delta signaling constrains reengineering of pro-T cells by PU.1. Proc Natl Acad Sci U S A 103, 11993–11998 (2006).

76. Taghon, T., Yui, M.A. & Rothenberg, E.V. Mast cell lineage diversion of T lineage precursors by the essential T cell transcription factor GATA-3. Nat Immunol 8, 845–55 (2007).

77. Yui, M.A., Feng, N. & Rothenberg, E.V. Fine-scale staging of T cell lineage commitment in adult mouse thymus. J Immunol 185, 284–293 (2010).

78. Meers, M.P., Bryson, T.D., Henikoff, J.G. & Henikoff, S. Improved CUT&RUN chromatin profiling tools. Elife 8 (2019).

79. Levanon, D., Negreanu, V., Lotem, J., Bone, K.R., Brenner, O., Leshkowitz, D. & Groner, Y. Transcription factor Runx3 regulates interleukin-15-dependent natural killer cell activation. Mol Cell Biol 34, 1158–69 (2014).

80. Liu, N., Hargreaves, V.V., Zhu, Q., Kurland, J.V., Hong, J., Kim, W., Sher, F., Macias-Trevino, C., Rogers, J.M., Kurita, R. et al. Direct Promoter Repression by BCL11A Controls the Fetal to Adult Hemoglobin Switch. Cell 173, 430–442 e17 (2018).

81. Langmead, B. & Salzberg, S.L. Fast gapped-read alignment with Bowtie 2. Nat Methods 9, 357–9 (2012).

82. Heinz, S., Benner, C., Spann, N., Bertolino, E., Lin, Y.C., Laslo, P., Cheng, J.X., Murre, C., Singh, H. & Glass, C.K. Simple combinations of lineage-determining transcription factors prime cis-regulatory elements required for macrophage and B cell identities. Mol Cell 38, 576–89 (2010).

83. McLean, C.Y., Bristor, D., Hiller, M., Clarke, S.L., Schaar, B.T., Lowe, C.B., Wenger, A.M. & Bejerano, G. GREAT improves functional interpretation of cis-regulatory regions. Nat Biotechnol 28, 495–501 (2010).

84. Ramirez, F., Ryan, D.P., Gruning, B., Bhardwaj, V., Kilpert, F., Richter, A.S., Heyne, S., Dundar, F. & Manke, T. deepTools2: a next generation web server for deep-sequencing data analysis. Nucleic Acids Res 44, W160–5 (2016).

85. Hao, Y., Hao, S., Andersen-Nissen, E., Mauck, W.M., 3rd, Zheng, S., Butler, A., Lee, M.J., Wilk, A.J., Darby, C., Zager, M. et al. Integrated analysis of multimodal single-cell data. Cell 184, 3573–3587 e29 (2021).

86. Qiu, X., Mao, Q., Tang, Y., Wang, L., Chawla, R., Pliner, H.A. & Trapnell, C. Reversed graph embedding resolves complex single-cell trajectories. Nat Methods 14, 979–982 (2017).

87. Trapnell, C., Cacchiarelli, D., Grimsby, J., Pokharel, P., Li, S., Morse, M., Lennon, N.J., Livak, K.J., Mikkelsen, T.S. & Rinn, J.L. The dynamics and regulators of cell fate decisions are revealed by pseudotemporal ordering of single cells. Nat Biotechnol 32, 381–386 (2014).

