## Supplementary_Figures for "Runx factors launch T-cell and innate lymphoid programs via direct and gene network-based mechanisms"

### Supplementary Figure 1

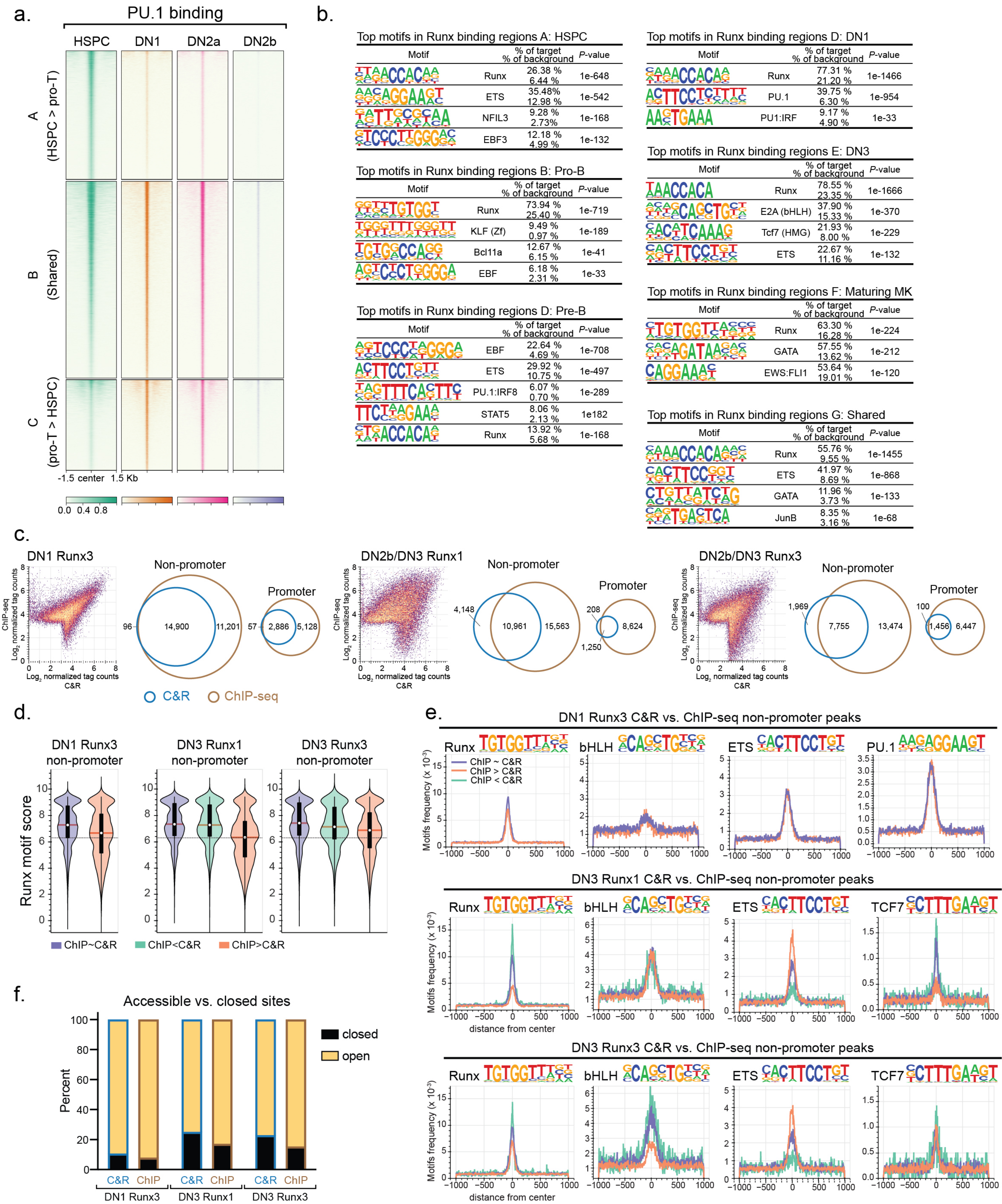

### Supplementary Figure 2

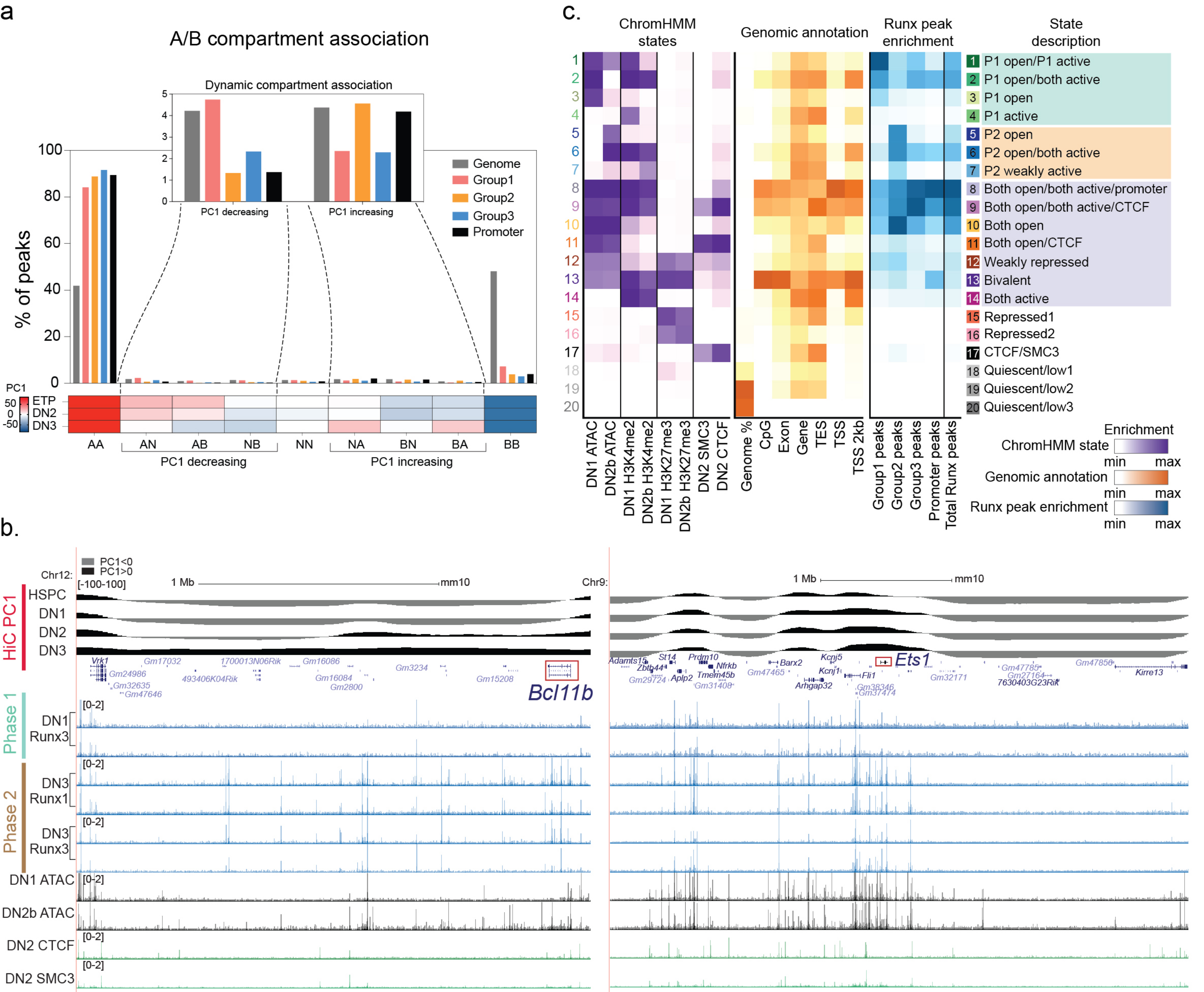

Supplementary Figure 3

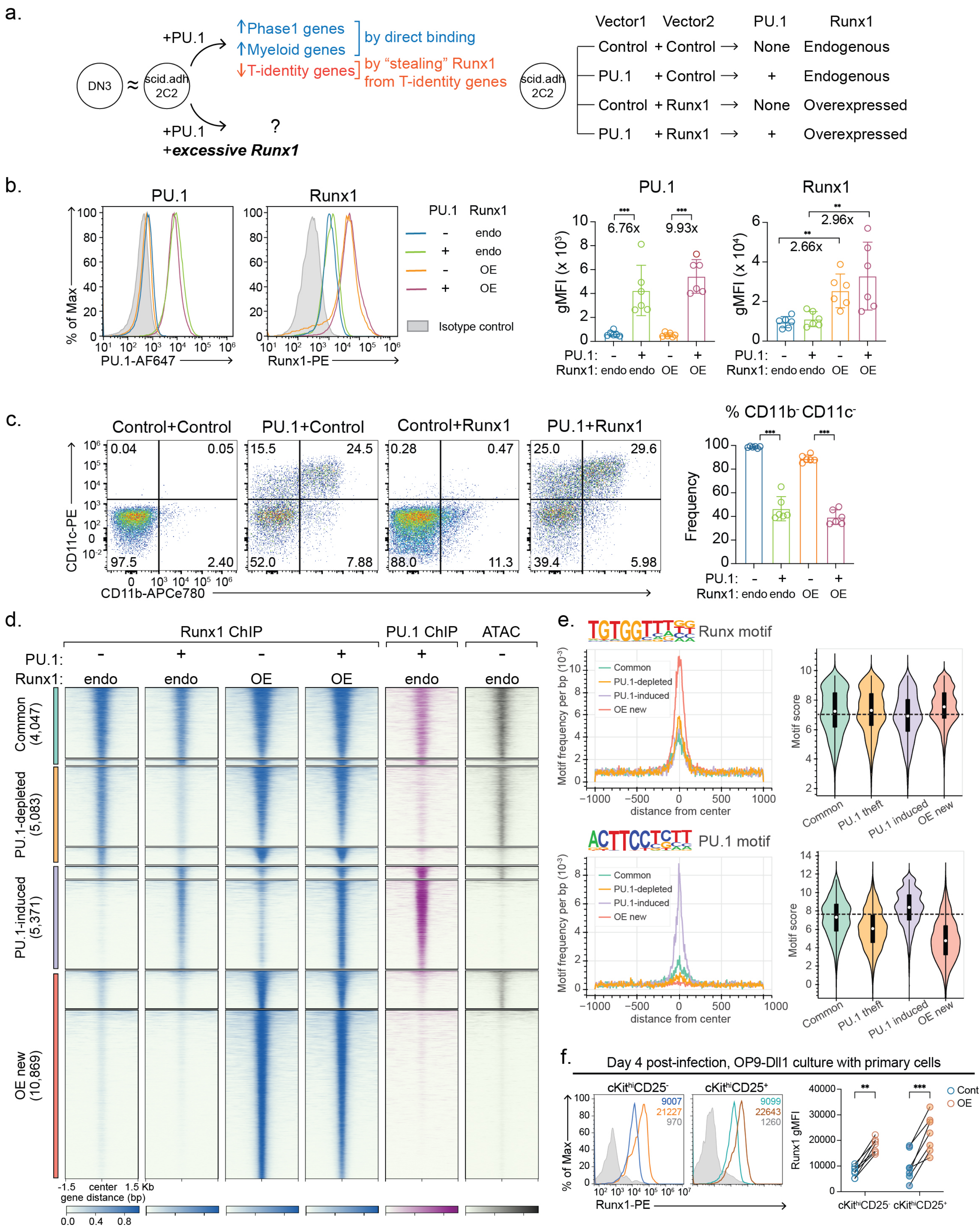

### Supplementary Figure 4

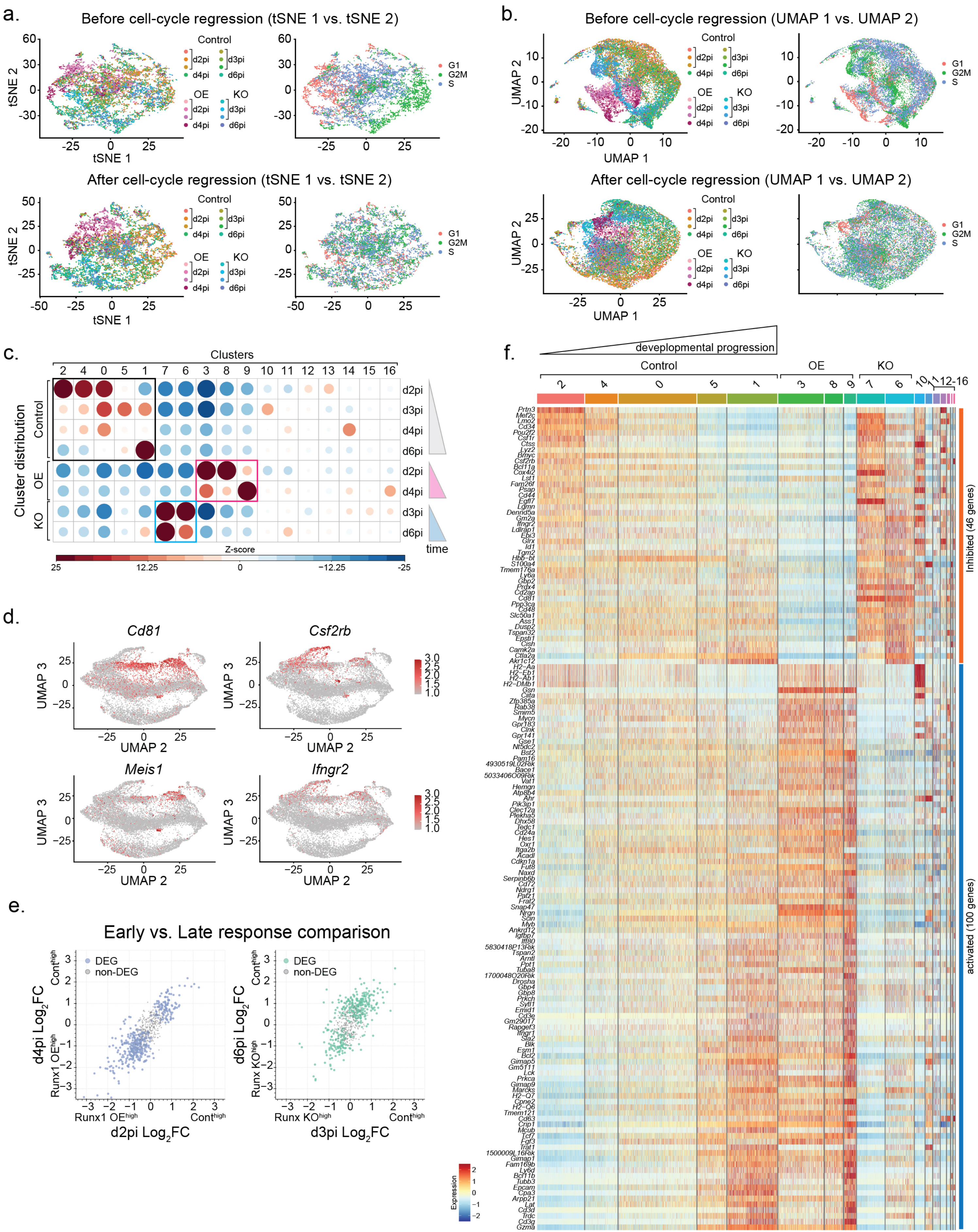

### Supplementary Figure 5

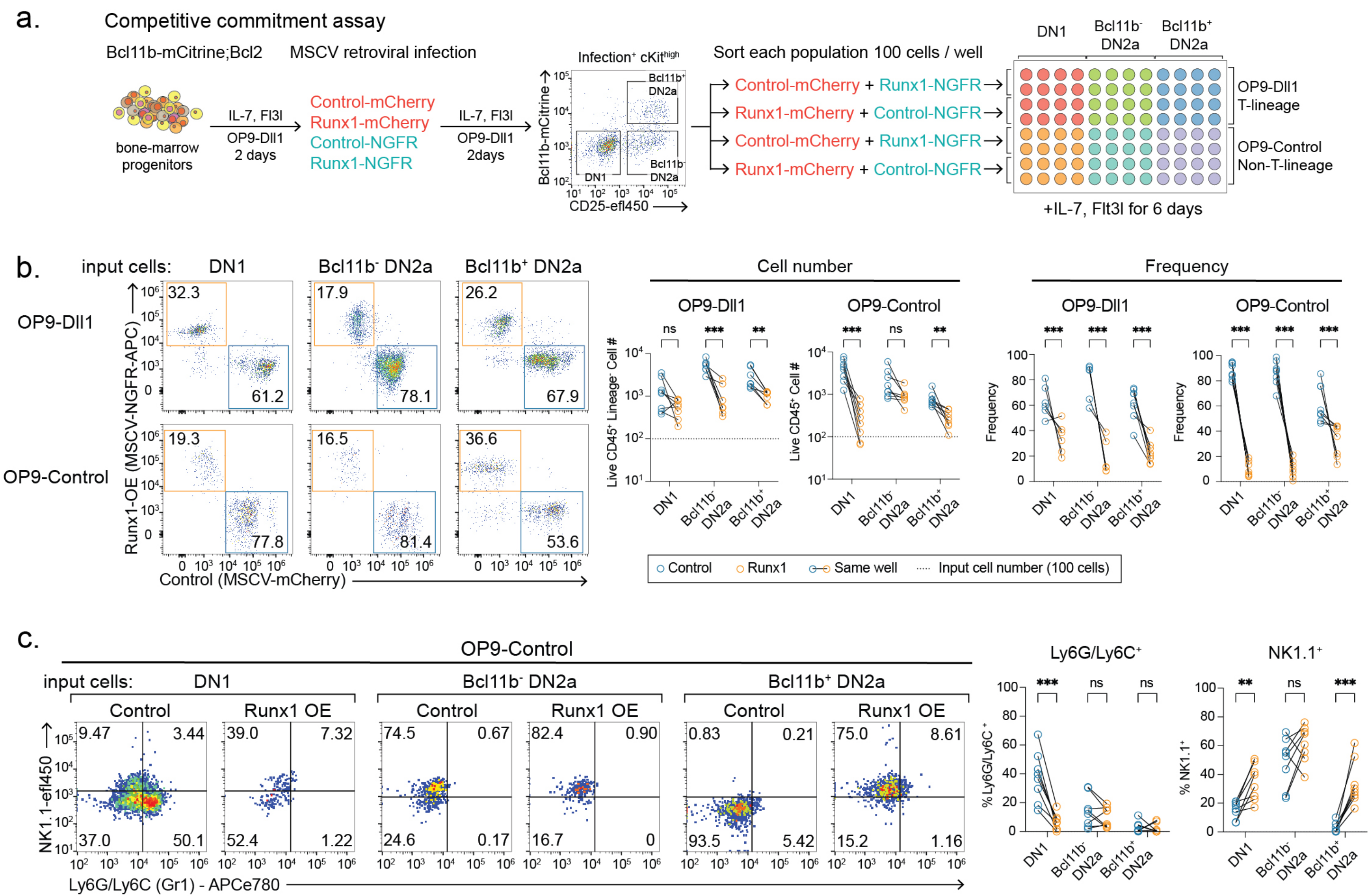

### Supplementary Figure 6

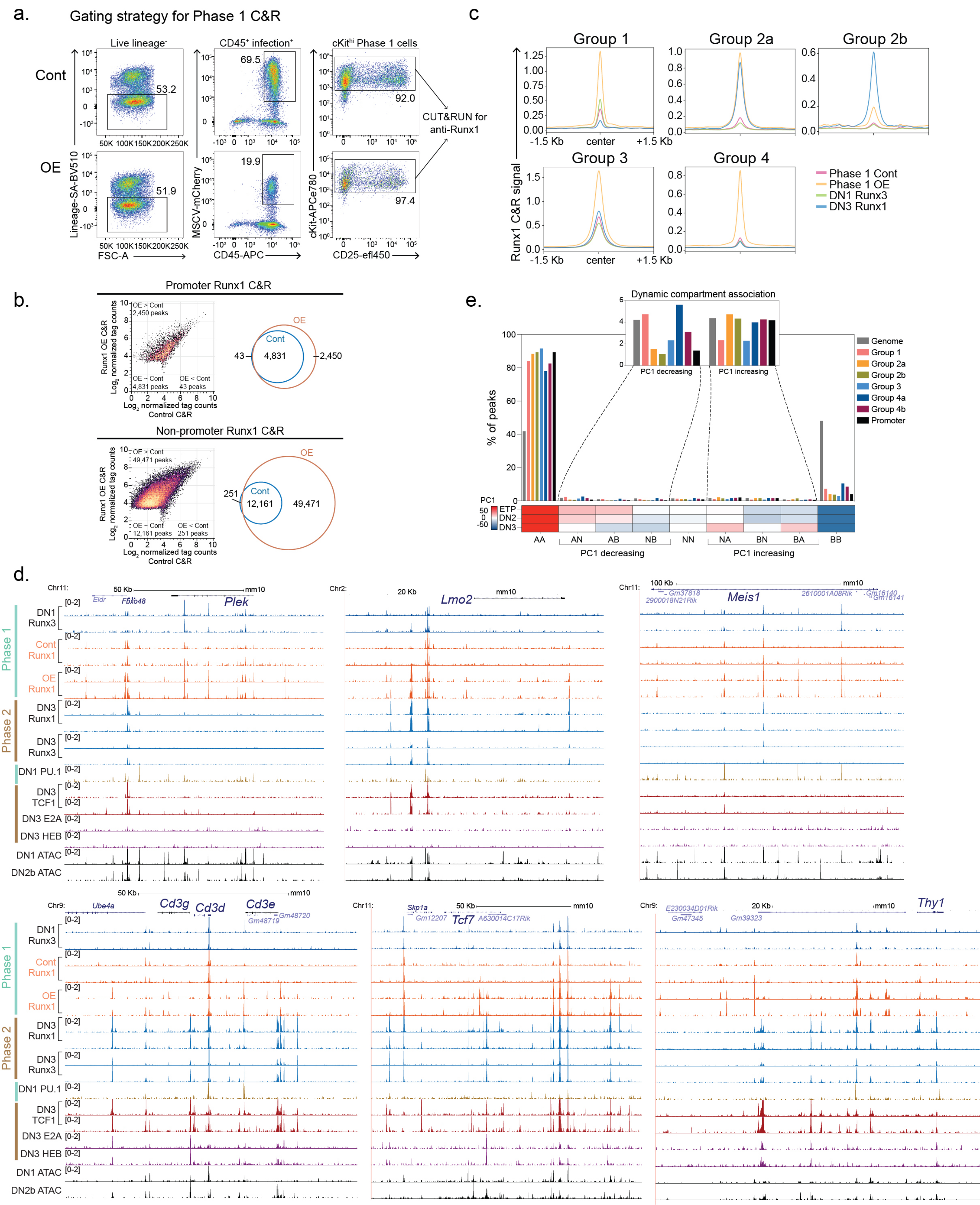

### Supplementary Figure 7

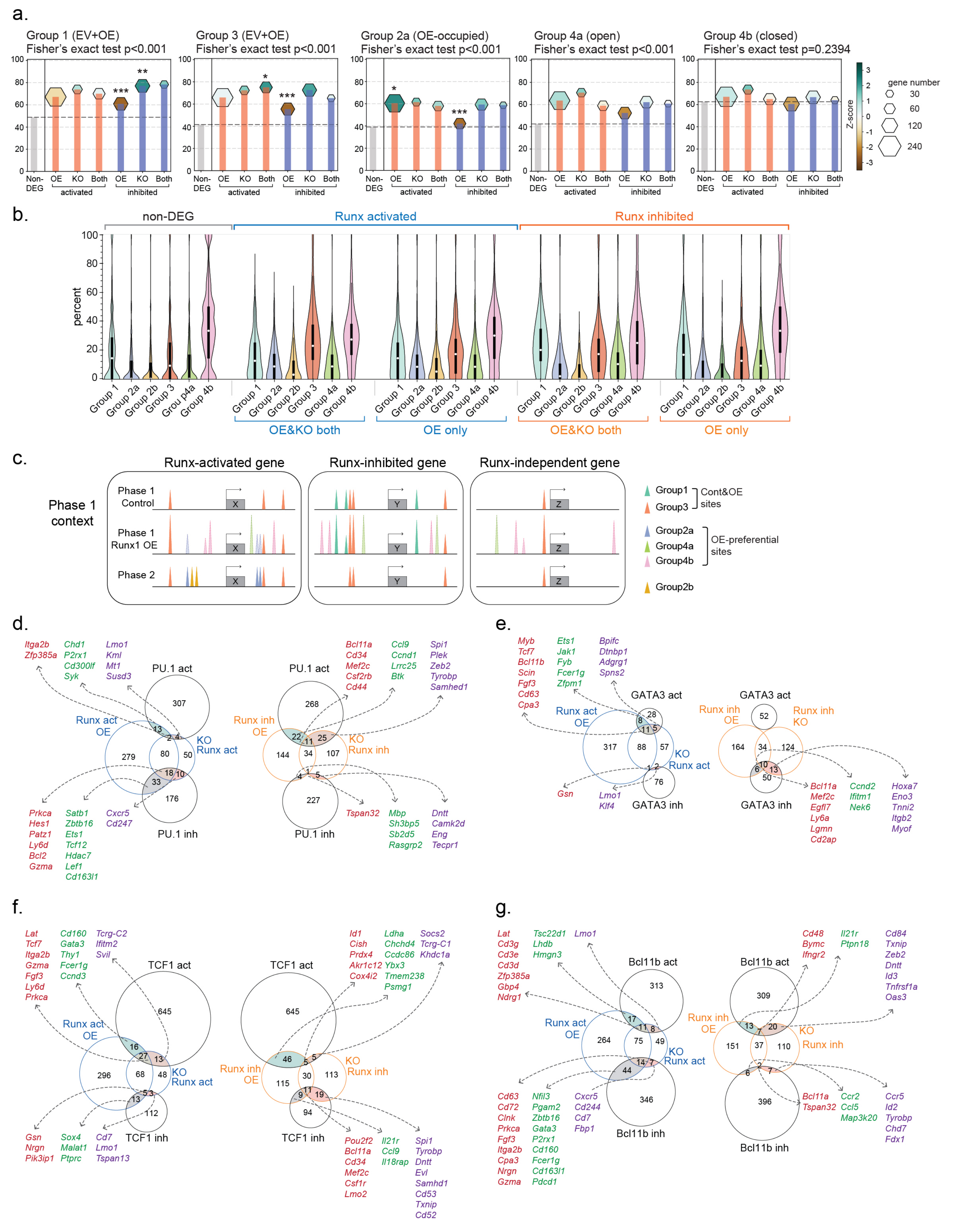
